# Cryo-EM structure of an activated GPCR-G protein complex in lipid nanodiscs

**DOI:** 10.1101/2020.06.11.145912

**Authors:** Meng Zhang, Miao Gui, Zi-Fu Wang, Christoph Gorgulla, James J Yu, Hao Wu, Zhen-yu Sun, Christoph Klenk, Lisa Merklinger, Lena Morstein, Franz Hagn, Andreas Plückthun, Alan Brown, Mahmoud L Nasr, Gerhard Wagner

**Affiliations:** Department of Biological Chemistry and Molecular Pharmacology, Blavatnik Institute, Harvard Medical School, Boston, MA, USA; Center for Integrated Protein Science Munich (CIPSM) at the Department of Chemistry and Institute for Advanced Study, Technical University of Munich, Garching, Germany; Department of Biochemistry, University of Zurich, Zurich, Switzerland; Department of Medicine, Brigham and Women’s Hospital, Harvard Medical School, Boston, MA, USA

## Abstract

G protein coupled receptors (GPCRs) are the largest superfamily of transmembrane proteins and the targets of over 30% of currently marketed pharmaceuticals^1,2^. Although several structures have been solved for GPCR-G protein complexes^3–17^, structural studies of the complex in a physiological lipid membrane environment are lacking. Additionally, most previous studies required additional antibodies/nanobodies and/or engineered G proteins for complex stabilization. In the absence of a native complex structure, the underlying mechanism of G protein activation leading to GDP/GTP exchange remains unclear. Here, we report cryo-EM structures of lipid bilayer-bound complexes of neurotensin, neurotensin receptor 1, and Gα_i1_β_1_γ_1_ protein in two conformational states, resolved to 4.1 and 4.2 Å resolution. The structures were determined without any stabilizing antibodies/nanobodies, and thus provide a native-like platform for understanding the structural basis of GPCR-G protein complex formation. Our structures reveal an extended network of protein-protein interactions at the GPCR-G protein interface compared to in detergent micelles, defining roles for the lipid membrane in modulating the structure and dynamics of complex formation, and providing a molecular explanation for the stronger interaction between GPCR and G protein in lipid bilayers. We propose a detailed allosteric mechanism for GDP release, providing new insights into the activation of G proteins for downstream signaling under near native conditions.

---

G protein coupled receptors (GPCRs) sense extracellular stimuli including odorants, hormones, neurotransmitters, and photons. A stimulus leads to a shift in the conformational equilibrium of the GPCR towards a state which favors binding of the intracellular signal transducer, GDP-bound heterotrimeric Gαβγ protein^18^. Binding causes perturbation of the GDP binding pocket, leading to replacement of GDP by GTP and the dissociation of the Gα and Gβγ subunits from each other and the GPCR^19^. The released Gα and Gβγ subunits remain anchored to the membrane through lipid modifications but diffuse and interact with downstream effectors to stimulate signaling cascades^18^.

Recent advances in X-ray crystallography and cryo-EM have allowed the determination of several GPCR-G protein complex structures to near-atomic resolution^3–8,11–17,20^. However, due to difficulties in preparing stable GPCR-G protein complexes in detergent micelles, a range of stabilization techniques had to be employed that compromise the function of the complexes. These include: i) antibodies or nanobodies that prevent GTP binding to Gα; ii) engineered dominant-negative Gα (DNGα) subunits that do not exhibit GDP/GTP exchange activity; or iii) engineered mini-G proteins that lack the α-helical domain (AHD) of Gα. Furthermore, all the previous structural studies reconstituted GPCR-G protein complexes in detergent micelles, which fail to replicate the properties of the native lipid bilayer environment of GPCRs, including membrane thickness, lateral pressure, and curvature^21^. It has been reported that various GPCRs exhibit higher stability and better functionality when incorporated into lipid bilayers as compared to detergent micelles^22,23^. Additionally, negatively charged lipids have been found to allosterically modulate GPCR activation and its selective interaction with G proteins^24–26^. Therefore, structural and dynamical information for the GPCR–G protein interaction in a lipid bilayer environment is necessary to understand the GPCR signal transduction mechanism.

To investigate the interaction between GPCR and G proteins in lipid bilayers, we used the neurotensin receptor 1 (NTR1)-G_i_ interaction as a model system. NTR1 is a class A GPCR that responds to neurotensin (NT), a 13-residue peptide implicated in the pathogenesis of schizophrenia, antinociception, hypothermia, Parkinson’s disease and tumor growth^1,27^. To reconstitute and determine the structure of the NT_8-13_-NTR1-Gα_i1_β_1_γ_1_ complex in a lipid bilayer environment we used circularized nanodiscs (cNDs) prepared with covalently circularized membrane scaffold proteins^28^. While structures of the GDP-bound G protein heterotrimer^29^ and GPCR-G protein complexes were so far only available in detergent micelles, such as the recent cryo-EM structure of the NTR1-Gα_i1_β_1_γ_2_ complex stabilized by an antibody (scFv16) and in complex with a pseudopeptide analog of NT^8^, the data shown here provide insights into the mechanism by which a G protein is activated by the interaction with GPCR in a lipid bilayer.

## Lipid bilayers enhance the efficiency of NT-NTR1-Gα_i1_β_1_γ_1_ complex formation

To enable efficient expression of NTR1 for purification and structural studies, we took advantage of the TM86V-L167R ΔIC3B construct^30^. Among all NTR1 evolved variants, TM86V-L167R ΔIC3B shows most wild-type-like GDP/GTP exchange activities^30^ and exhibits similar downstream signaling functionality to wild-type NTR1 as measured by the production of inositol-1-phosphate (IP1), the final metabolite of the inositol phosphate cascade, with a EC50 of 2.7 nM for wild-type NTR1 and 0.22 nM for TM86V-L167R ΔIC3B (Extended Data Fig. 1a, left). Interestingly, a single mutation of R167^3.50^L (superscripts denote Ballesteros–Weinstein numbering^31^) completely quenched IP1 production (Extended Data Fig. 1a, right). As we discuss later in details, R167^3.50^ is involved in direct interaction with G_i_ in the complex structures determined in this study, explaining partially the critical impact of this residue in the signaling process.

NTR1 was affinity purified using immobilized NT_8-13_, which ensured selection of properly folded NTR1 only. The purified NT-NTR1 complex was then incorporated into 9-nm diameter covalently circularized nanodiscs (cNDs), containing a mixture of zwitterionic lipid POPC and negatively charged lipid POPG, and belted by circularized membrane scaffold protein cNW9 (ref.^28^) (Fig. 1a and Extended Data Fig. 1). Heat-treating the purified nanodiscs at 42 °C for 24 hours improved sample homogeneity (Extended Data Fig. 1c). Circular dichroism measurements showed increased thermostability of NTR1 in cNDs as compared to in detergent micelles, with a transition temperature about 18 °C higher (Fig. 1b and Extended Data Fig. 2a, b), implying different dynamic properties in lipid bilayer relative to in detergents. The sample was stable at 45 °C for at least 15 days, showing well dispersed and reproducible peaks on two-dimensional nuclear magnetic resonance (2D NMR) spectra (Extended Data Fig. 3a). These observations agree with studies showing that GPCRs are more stable in membrane environments^32^. When Gα_i1_β_1_γ_1_ was incorporated into cNDs using the same method, its thermostability also improved relative to in detergent micelles (Extended Data Fig. 2d-f).

**Fig. 1.**
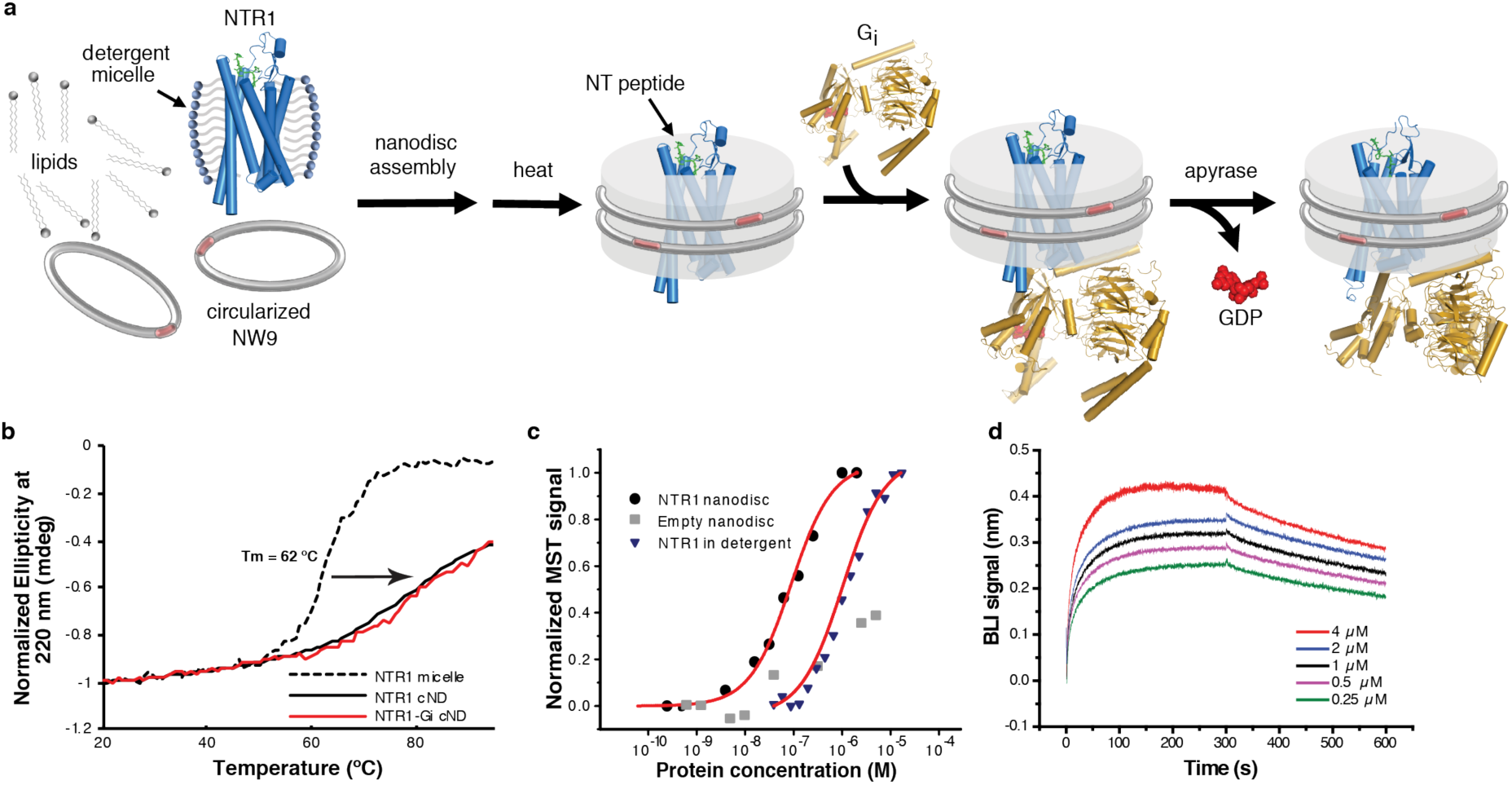
Assembly and biophysical characterization of the NT-NTR1-Gα_i1_β_1_γ_1_ complex in circularized nanodiscs (cNDs). **a**, Schematic showing the assembly of the NT-NTR1-G_i_ complex in lipid nanodiscs using the circularized membrane scaffold protein cNW9. **b**, Circular dichroism thermostability assays on NT-NTR1 in detergent micelles (dashed black line) with NT-NTR1 (solid black line) and NT-NTR1-G_i_ complex (solid red line) in cNDs. **c**, Microscale thermophoresis data fitting for the interaction between NT-NTR1 and G_i_ in diheptanoylphosphatidylcholine detergent (DH_7_PC) yields a K_D_ of 1400±100 nM (blue triangles). The interaction between NT-NTR1-cND and G_i_ (black circles) yields a K_D_ of 76±18 nM. Weak binding between empty nanodiscs and G_i_ is shown as gray squares. **d**, Bio-layer interferometry (BLI) traces of G_i_ binding to NT-NTR1-cND at five different concentrations. Data fitting results are shown in Extended Data Fig. 4a, b.

To reconstitute the physiological signaling complex, we incubated NT-NTR1-cND with wild-type heterotrimeric human Gα_i1_β_1_γ_1_, which is myristoylated on Gα_i1_ and prenylated on Gγ_1_ (Extended Data Fig. 1d). The NT-NTR1-Gα_i1_β_1_γ_1_ complex in cNDs exhibits high thermostability (Fig. 1b and Extended Data Fig. 2c), and the binding affinity of NTR1 to Gα_i1_β_1_γ_1_ is higher in cNDs than in detergent micelles (K_D_ of 76 nM compared to 1.4 μM) (Fig. 1c), reflecting the essential role the membrane plays in efficient GPCR-G protein complex formation. Further binding kinetic measurements revealed two binding modes in cNDs with K_D_ of 5.8 nM and 38 nM, respectively (Fig. 1d and Extended Data Fig. 4a, b). The complex in cND is capable of GDP/GTP exchange, as shown by a much higher dissociation rate upon addition of GTPγS (Extended Data Fig. 4c). However, for the following structural studies, we used apyrase to hydrolyze free GDP, which stabilizes the NT-NTR1-Gα_i1_β_1_γ_1_ complex.

## Cryo-EM structure of the NT-NTR1-Gα_i1_β_1_γ_1_ complex in cNDs

The higher affinity and improved thermostability of the NT-NTR1-Gα_i1_β_1_γ_1_ complex in lipid bilayers relative to in detergent micelles allowed us to collect cryo-EM data (Fig. 2, Extended Data Fig. 5) for the complexes without the need for further stabilization by antibodies/nanobodies or engineered G proteins. Two-dimensional class averages showed intact complexes within cNDs with uniform 9-nm diameters (Extended Data Fig. 5). Three-dimensional classification of these projections revealed two well-resolved classes, corresponding to “canonical” (C) and “noncanonical” (NC) states of the NT-NTR1-Gα_i1_β_1_γ_1_ complex, at 4.3 and 4.5 Å resolution, respectively (Extended Data Fig. 5). Two conformational states were also seen in the recent cryo-EM study of the scFv16-stabilized NTR1-Gα_i1_β_1_γ_2_ complex in detergent micelles^8^, but, as we describe below, these states are different from those that we observe (Fig. 2c). Additional density surrounds NTR1, corresponding to the cNW9 membrane scaffold protein and the lipid bilayer it encloses. Masking out these densities improved the resolutions of the C and NC states to 4.1 Å and 4.2 Å respectively (Extended Data Fig. 5). In these maps, the pitch of helices and many sidechains are clearly resolved (Extended Data Fig. 6), allowing us to confidently place and remodel known atomic models of NT, NTR1^30^ and Gα_i1_β_1_γ_1_ (ref. ^9,30^). The density of NT is well revolved in both conformations (Extended Data Fig. 6), and adopts similar structure and interactions to those observed in detergent micelles^8,33^. The N-terminal helices of Gβ and Gγ both show weak densities, presumably due to flexibility.

**Fig. 2.**
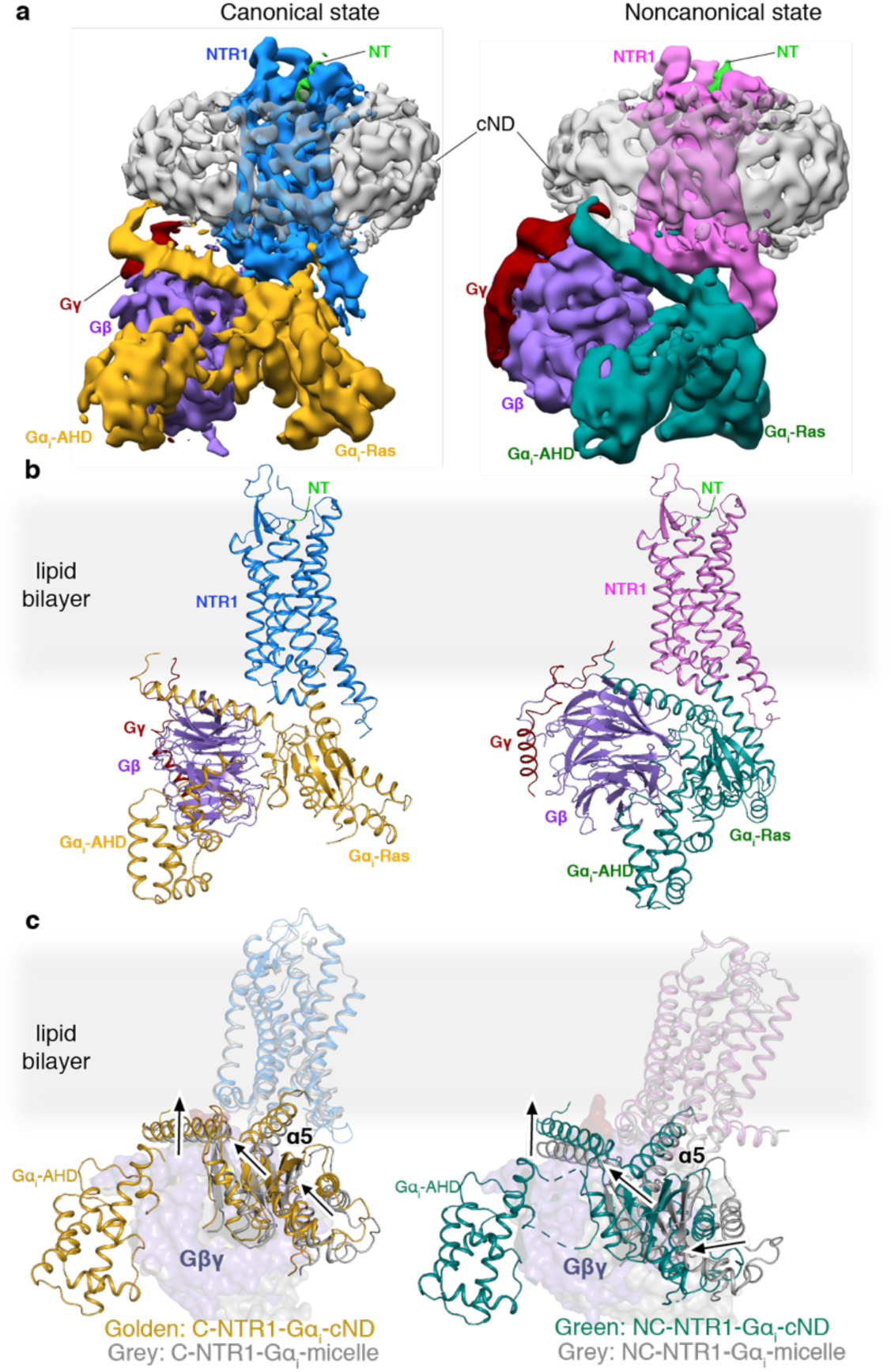
Cryo-EM structures of NT-NTR1-G_i_-cND. **a**, Cryo-EM density maps of NT-NTR1-G_i_-cND complex in the canonical state (left) and in the noncanonical state (right). The maps are low-pass filtered to 5 Å and colored by subunit. Higher-resolution maps were obtained by masking out density for the nanodisc and Gα−AHD domain. **b**, Atomic models of NT-NTR1-G_i_-cND complex in the canonical state (left) and in the noncanonical state (right). The models are shown in the same orientation as the maps in (**a**). **c**, Structural superimposition of C-NT-NTR1-G_i_-cND with C-NTR1-scFv16-micelle (left) and NC-NT-NTR1-G_i_-cND with NC-NTR1-scFv16-micelle (right). Structural displacement is highlighted with arrows. The models are superimposed on the NTR1.

Compared to most reported structures ^3,4,6,8,11–14,16,17^, the α-helical domain (AHD) of Gα_i1_ is clearly resolved in both states (Fig. 2a, Extended Data Fig. 7a, b). In the few structures that do report the position of the AHD^5,15,20^, the position may be affected by crystal contacts and/or the nanobodies/antibodies that were included for stabilization (Extended Data Fig. 7c-f). Our structures lack these constraints and therefore more closely reflect the native orientation and localization of the AHD in the nucleotide-free state. In comparison to the crystal structure of the GDP-bound G_i_ trimer^29^, the AHD moves away from its close association with the Ras-like domain of Gα and interacts with the outer strands of the second and third β blades of Gβ after GDP release (Fig. 2b, Extended Data Fig. 7a-c). As we discuss later, the large-scale movement of AHD is an important step in the GDP release pathway.

## Lipid bilayer modulates GPCR-G protein interaction

The NT-NTR1-Gα_i1_β_1_γ_1_ complex shows clear interactions with the lipid bilayer in both the C and NC states (Fig. 3a, Extended data Fig. 8). Density at the beginning of the αN-helix of Gα is observed protruding into the lipid bilayer, which corresponds to the myristoylation site of the Gα (Fig. 3b, top panel). Similar density at the C-terminus of Gγ corresponds to the prenylation site (Fig. 3b, top panel). These lipid moieties anchor the G protein to the membrane. Lipid density is also observed above the positively charged αN-helix of Gα (Fig. 3b, bottom panel). The sidechains of arginine and lysine residues within this helix are oriented towards the membrane and likely form electrostatic interactions with the negatively charged lipid POPG (Fig. 3b, bottom panel). This observation agrees with the finding that negatively charged lipids strengthen the interaction between NTR1 and G protein^26^. In comparison with other structures of class A GPCR-G_i_ complexes, the αN-helices of C and NC states solved here are located close to the membrane, while they bend away in structures determined in detergent micelles (Fig. 3c). The observed hydrophobic and electrostatic interactions ensure close proximity of G_i_ to NTR1, and thus enhance G_i_ binding to NTR1, particularly between the αN-β1 hinge of G_i_ and ICL2 of NTR1 as described below (Fig. 4a).

**Fig. 3.**
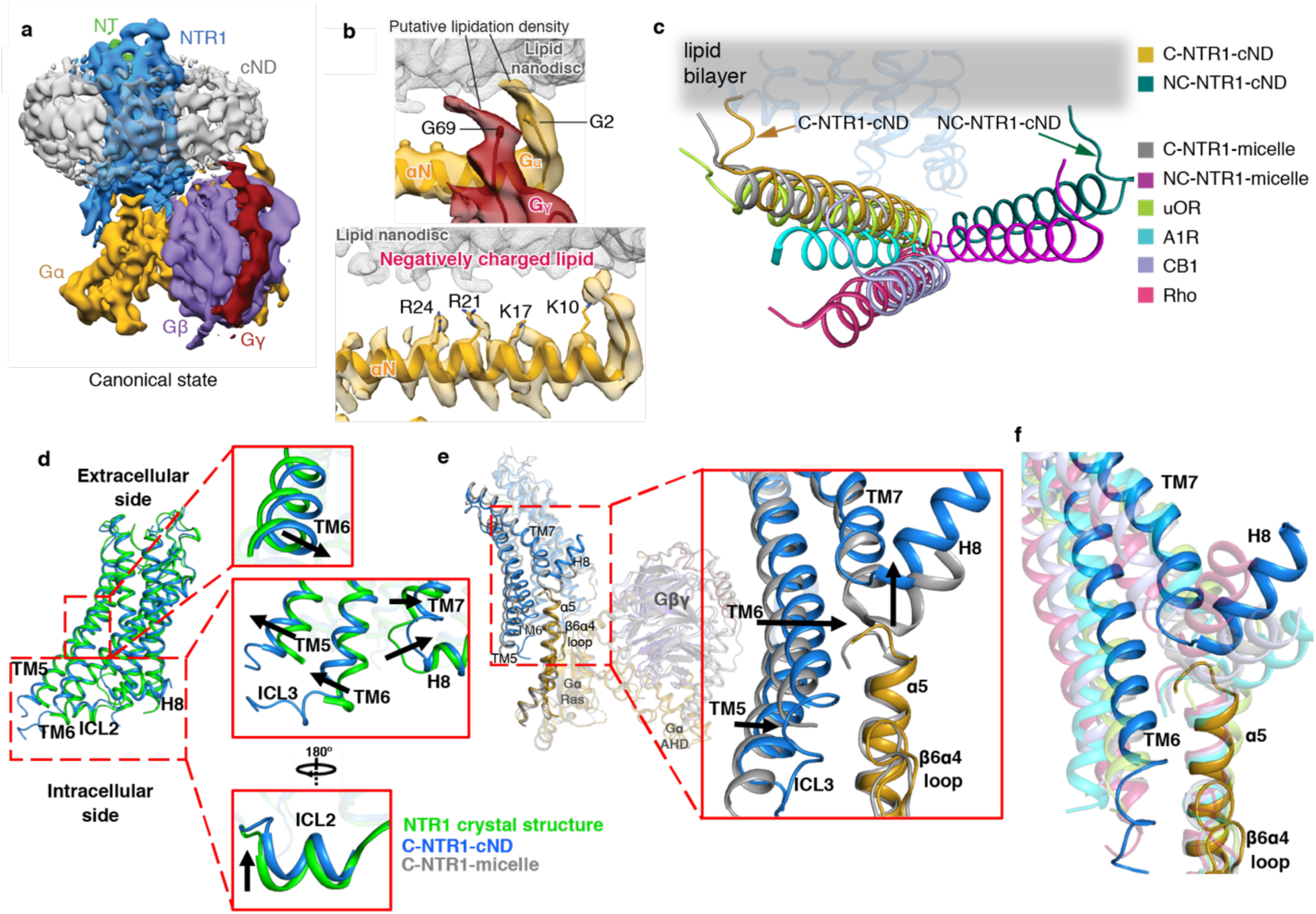
Impact of lipid bilayer on the NTR1-G_i_ complex. **a**, Cryo-EM density map of the NT-NTR1-G_i_-cND complex in the canonical state. The map is low-pass filtered to 5 Å and colored by subunit. **b**, Top panel, density for the putative lipid modifications of glycine 2 (G2) of Gα_i1_ and glycine 69 (G69) of Gγ_1_. Nanodisc density is shown as gray mesh. The density map of the canonical state is low-pass filtered to 5 Å. Bottom panel, positively charged residues of the αN helix of Gα_i1_ face the negatively charged lipid bilayer. The 4.1 Å density map of the canonical state is shown. **c**, Comparison of the αN helices of GPCR-G_i_ complexes. C-NTR1-cND and NC-NTR1-cND indicate the canonical and noncanonical states of the NT-NTR1-G_i_ complex in lipid nanodiscs. C-NTR1-micelle and NC-NTR1-micelle indicate the canonical and noncanonical states of JMV449-NTR1-G_i_ complex in detergent micelles. Other Class A GPCR-G_i_ complexes: μOR-G_i_ (lime green; PDB 6DDE), A_1_R-G_i_ (cyan; PDB 6D9H), CB1-G_i_ (purple; PDB 6N4B), and Rho-G_i_ (hot pink; PDB 6CMO). The models are superposed on the GPCR. **d**, Structural comparison between NTR1 from the canonical state NT-NTR1-G_i_ complex in lipid nanodiscs (blue) and the crystal structure of NTR1 in detergent (green). Zoomed-in views are shown on the right. **e**, Structural comparison between the canonical states of NTR1-G_i_ in lipid bilayer (blue) and detergent (gray), superposed on the Ras-like domain of Gα (gold). Zoomed-in view of the cytoplasmic side of TM5-TM6, ICL3, TM7-H8, as well as the α5 helix and β6α4 loop of Gα is shown on the right. **f**, Comparison of the location of TM6 relative to the α5 helix of Gα in the canonical state NTR1 (blue) in complex with G_i_ (gold) with other class A GPCR-G_i_ complex structures, including the canonical state of NTR1-G_i_ in detergent micelle (gray), μOR-G_i_ (lime green), Rho-G_i_ (hot pink), A_1_R-G_i_ (cyan), and CB1-G_i_ (purple). The models are superposed on the Ras-like domain of Gα.

**Fig. 4.**
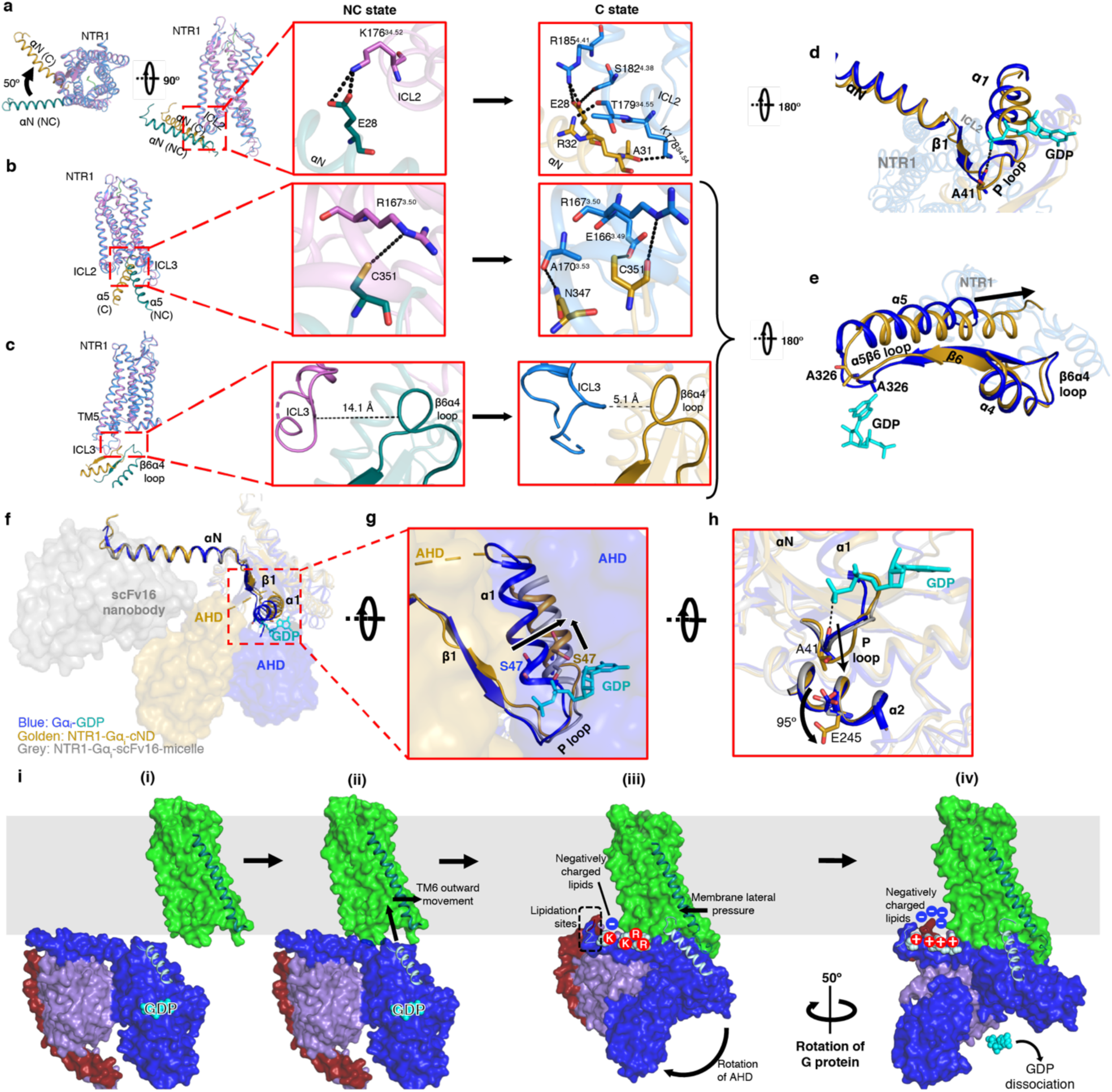
Allosteric modulation of the GDP binding pocket by the NTR1-G_i_ interaction. **a-c**, Superposition between C-state NTR1 (blue) and αN helix of Gα (gold) with the NC-state NTR1 (orchid) and αN helix of Gα (dark cyan). Models are superposed on NTR1. Overview (left) and zoomed-in views of the NC state (middle) and C state (right) are shown. **a**, ICL2-αN helix interactions. Compared to the NC state, the αN helix of Gα of the C state is rotated by 50°. **b**, NTR1-α5 helix interactions. **c**, ICL3-β6α4 loop interactions. The backbones of ICL3 and β6α4 are closer in the C state and form interactions predicted by molecular dynamics simulations (Extended Data Fig. 11a). **d**, Intracellular view showing perturbation of the P loop in the C state (gold) relative to the crystal structure of GDP-bound G_i_ (blue; GDP in cyan). **e**, Intracellular view showing perturbation of the α5β6 loop in the C state (gold) relative to the crystal structure of GDP-bound G_i_ (blue; GDP in cyan). In d-e, the models are superposed on the Gα Ras-like domain. **f**, Structures of GDP-bound Gα_i_ (blue; GDP in cyan), NTR1-bound Gα_i_ in detergent (grey) and NTR1-bound Gα_i_ in lipid bilayer (gold) showing the different locations of the AHD and the stabilizing antibody scFv16. The structures are superposed on αN-β1. **g**, Zoom-in view showing lateral displacement of α1 helix including S47 from the phosphates of GDP in NTR1-G_i_-cND. **h**, Rotation of E245 in NT-NTR1-G_i_-cND (gold) by 95° compared to the GDP-G_i_ structure (blue) to accommodate the P-loop. This structural change is not observed in detergent (grey). **i**, Model of the proposed insertion-rotation mechanism: (i) Lateral diffusion of NTR1 and G_i_ in the membrane; (ii) Recognition of NTR1 by G_i_, allowing insertion of α5 into the open cavity of NTR1; (iii) Formation of the NC state including displacement of the AHD; (iv) Formation of the C state following rotation of G_i_.

As expected, the majority of NTR1 is buried inside the lipid bilayer, including TM1-4 and TM7, the N-terminal half of TM5, and the C-terminal half of TM6. ICL2 and H8 are partially buried at the membrane surface (Extended Data Fig. 8c). To reveal the effects of the lipid bilayer on the GPCR, we compared our structures with the crystal structure of NTR1 (X-rNTR1)^33^ and the cryo-EM structure of NTR1 in the canonical state (C-hNTR1)^8^ (representing structures of agonist-bound NTR1 in detergent in the absence and presence of G_i_, respectively). In lipid bilayers, the core of NTR1 is more compact due to an inward movement of the middle of TM6 (Fig. 3d, Extended Data Fig. 9a), whereas X-rNTR1 and C-hNTR1 superpose well with each other (Extended Data Fig. 9b). Compression of TM6 is likely due to lateral pressure from the lipid bilayer, implying modulation of the folding and dynamics of NTR1 by the lipid bilayer. Relative to the detergent structures, ICL2 and the cytoplasmic side of TM7 and H8 show an upward movement, indicative of membrane association (Fig. 3d, Extended Data Fig. 9a). Overall, the increased compaction and better membrane association of NTR1 agrees with the improved thermostability observed in lipid bilayers (Fig. 1b, Extended Data Fig. 2c).

Upon insertion of the α5 helix of Gα into the core of NTR1, the cytoplasmic side of TM5, TM6 and ICL3 move outward to accommodate the α5 helix (Fig. 3d). Structural and dynamical changes are also observed in 2D NMR experiments on ^1^H^15^N-NTR1 upon binding to G_i_ in cNDs (Extended Data Fig. 3c). In the presence of the lipid bilayer, this movement appears to be more restricted than the large outward movement observed in detergent, potentially due to the lateral pressure from the lipid bilayer (Fig. 3e). The reduced movement of TM5 and TM6 relative to C-hNTR1 maintains closer contacts with the α5 helix (Fig. 3e). Comparison of TM6 positions among class A GPCR-G_i_ complexes reveals that TM6 in the C-state NTR1 exhibits closest proximity to the α5 helix, likely resulting in more potential interactions (Fig. 3f and Extended Data Fig. 9c). Taken together, these observations suggest that the lipid bilayer constrains the conformation of NTR1 to enhance its interaction with G_i_, agreeing with our observation of higher binding affinity in lipid bilayer (Fig. 1c).

## The NTR1-Gα_i1_β_1_γ_1_ interface

The C and NC states are related by a 50° rotation of G_i_ relative to NTR1 (Fig. 4a). This change in orientation results in different interactions between the αN helix and ICL2. In the C state, a potential salt bridge is observed between E28 and R185^4.41^, as well as several potential hydrogen bonds between E28 and S182^4.38^, R32 and T179^34.55^, and A31 and K178^34.54^ (Fig. 4a). In contrast, only one hydrogen bond (between R32 and T178^34.55^) is observed in C-hNTR1 in detergent micelles^8^. These additional contacts with ICL2 in the presence of the lipid bilayer likely result from the closer proximity of the αN helix to the membrane and NTR1 (Fig. 3c). Many of these interactions are absent in the NC state, where we observe only one potential salt bridge between E28 and K176^34.52^. Fewer contacts in the NC state suggest that it could be a less stable intermediate state before the final C-state complex.

The orientation of the α5 helix relative to NTR1 is also different between the two states, although the depth of insertion is the same (Fig. 4b). In the C state, several potential hydrogen bonds are observed, including C351 with E166^3.49^, C351 with R167^3.50^, and N347 with A170^3.53^ (Fig. 4b). The interaction between N347 and A170^3.53^ is also observed in C-hNTR1 ^8^. E166^3.49^ and R167^3.50^ belong to the highly conserved D/ERY motif. R167^3.50^ is found to be essential for downstream signaling (Extended Data Fig. 1a) and has been reported to be critical for GDP/GTP exchange through mutagenesis studies^30^. Examination of a range of class A GPCR-G_i_ structures shows that it is common for α5 insertion to stop at R^3.50^ (Extended Data Fig. 10d). Thus, R^3.50^ might serve as both an interaction hot-spot and an “access gate” that decides the depth of α5 insertion. The NC state displays fewer interactions with only one possible hydrogen bond between C351 and R167^3.50^ (Fig. 4b).

Rotation of G_i_ also results in the β6α4 loop moving closer to ICL3 in the C state than in either the NC state (Fig. 4c) or detergent structures (Extended Data Fig. 10b). Although the map quality of ICL3 prevents a detailed analysis, molecular dynamics simulations show potential salt bridges and hydrogen bonds forming between ICL3 and β6α4 loop in the C state (Extended Data Fig. 11). Similar interactions between ICL3 and the β6α4 loop have been observed in the structure of the adenosine A1 receptor (A_1_R)-Gα_i2_β_1_γ_2_ complex^12^.

## An insertion-rotation model for G_i_ activation

Comparison of our two conformational states with one another and with previous structures allows us to propose a mechanism of G-protein activation under near-native conditions. The presence of more GPCR-G_i_ contacts in the C state than the NC state, suggests that the NC state might be an intermediate, lower-affinity state. This implies that in addition to the close proximity between GPCR and G_i_ regulated by lipid bilayer, a certain orientation of G_i_ relative to GPCR is also required to enable efficient complex formation. This is consistent with our kinetics experiments which showed both high (5.8 nM) and lower affinity (38 nM) binding modes (Fig. 1d and Extended Data Fig. 4). A sequential model was also proposed to link the states observed with scFv16-stabilized hNTR1-G_i_ in detergent micelles^8^. Following this hypothesis, it appears that the interaction between NTR1 and G_i_ goes through an insertion-rotation mechanism (Fig. 4i). NTR1 and G_i_ first laterally diffuse in membrane until they encounter with each other. The cavity in NTR1 allows insertion of the α5 helix into the open core of NTR1. Subsequently, G_i_ rotates around α5 by approximately 50°, which maximizes protein-protein interactions (Fig. 4, Extended Data Fig. 10). The rotation stops when the β6α4 loop collides with ICL3, the αN-β1 hinge is caught by ICL2, and the α5 helix forms most contacts with the core of NTR1. As the α5 helix in the NT-NTR1-Gα_i1_β_1_γ_1_ C state exhibits one of the far-most rotated positions among class A GPCR-G_i_ complexes (Extended Data Fig. 10d), the NT-NTR1-Gα_i1_β_1_γ_1_ C state likely represents the final state enabling GDP dissociation (Fig. 4i).

## Multiple NTR1-G_i_ interactions stimulate the dissociation of GDP

Based on comparison of our structures with the structure of GDP-Gi^29^, we propose a multipartite mechanism for receptor-catalyzed nucleotide exchange (Fig. 5). In the unbound G-protein, the nucleotide is buried between the ras-homology domain (RHD) and the AHD of Gα. It has been suggested that when G protein encounters the receptor, α5 is straightened and forms early interactions with the GPCR, which initiates the GDP release process^34^. The AHD dissociates from the RHD, and, as we show here, interacts with the outermost strands of Gβ (Extended Data Fig. 7a, b). Previous computational simulations have shown that separation of the AHD is necessary (presumably to create an exit pathway for GDP) but not sufficient for rapid nucleotide release^35,36^. Here we observe that multiple allosteric pathways converge on structural rearrangement of the GDP binding site, and it is the combination of these pathways that are responsible GDP dissociation.

**Fig. 5.**
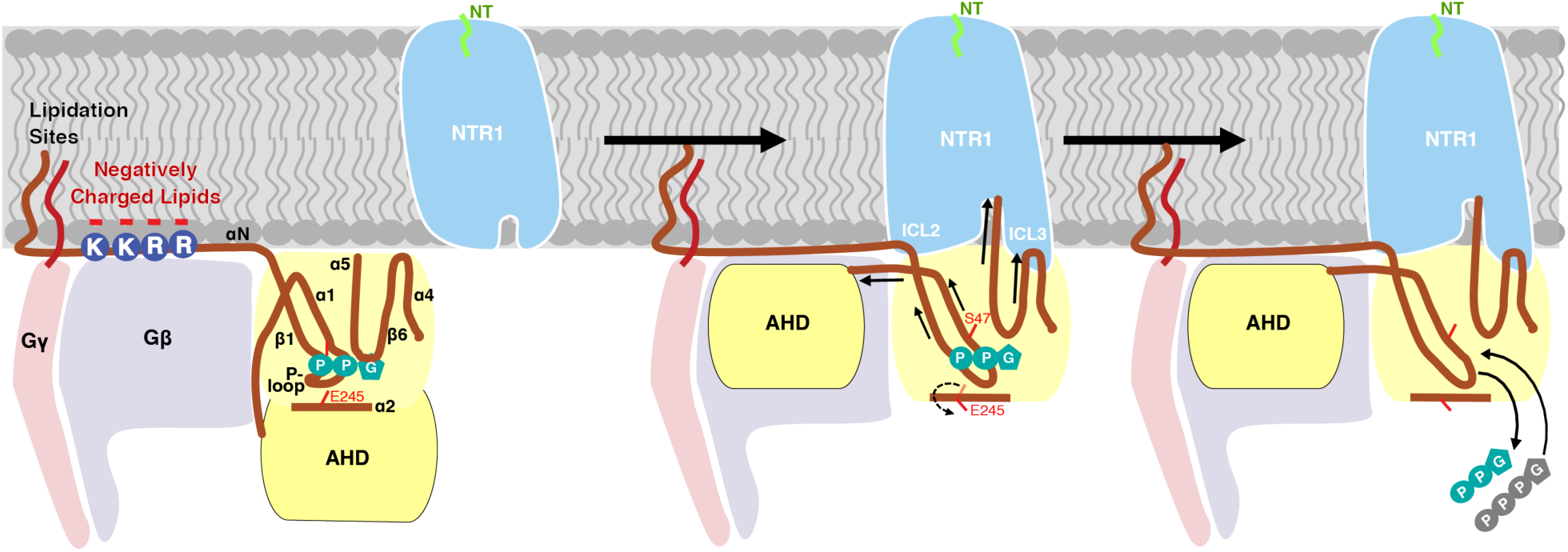
Proposed GDP release mechanism. The interaction between G_i_ and NTR1 leads to allosteric modulation of the GDP-binding site via three pathways: (1) The movement of the AHD to Gβ perturbs the directly linked α1, resulting in α1 dissociation from the phosphate groups of GDP; (2) The interaction between ICL2 of NTR1 and αN-β1 hinge of G protein delivers perturbation into the P-loop though β1, resulting in P-loop dissociation from the phosphate groups of GDP, which is coupled with a 95° rotation of the sidechain of E245 on α2; (3) The interactions between α5 and core of NTR1 and between β6α4 loop and ICL3 pull the α5β6 loop away from the guanine ring of GDP. The multi-point coordination of these structural elements leads to dissociation of both the phosphates and the guanine ring of GDP from G_i_, and thus dissociation of the entire GDP. Release of GDP creates a free nucleotide-binding pocket available for subsequent GTP binding, thus completing the GDP/GTP exchange process.

In the first pathway, insertion and rotation of the α5 helix into the core of NTR1 by two helical turns compared to the GDP-G_i_ structure^29^ displaces the α5β6 loop, which is responsible for binding the guanine ring of GDP in the nucleotide-bound state (Fig. 4e). This is consistent with previous structural studies showing that the α5β6 loop perturbation induced by the rotational translation of α5 helix is essential for GDP dissociation.^3,4,15–17,20,37,38,5,6,8,9,11–14^ As a result of this perturbation, A326 in the highly conserved TCAT motif moves away from its position in the GDP-G_i_ structure resulting in loss of contact with GDP. This agrees with previous mutagenesis studies showing that A326 is essential for GDP binding^39^. The conformation of the α5β6 loop is different from that in the detergent structure, potentially as a result of the different angles with which α5 inserts into NTR1 (Extended Data Fig. 10c, d). This agrees with computational simulations in which the tilt angle of the α5 helix was found to directly correlate with the conformation of the α5β6 loop^36^. The new conformation of the α5β6 loop, and therefore the dynamics of GDP loss, may be affected by the neighboring interaction between ICL3 and the β6α4 loop (Fig. 4c, e, Extended data Fig. 11).

In the second pathway, displacement of AHD appears to be correlated with movement of α1 (to which AHD is tethered) (Fig. 4f, g). This lateral movement causes residues within α1, including S47, to dissociate from the phosphate group of GDP (Fig. 4f, g). The S47N mutation is dominant negative^14^, suggesting that this movement is a key step towards GDP release. Our observation here agrees with previous mutagenesis^40^, HDX^37^ as well as computational^41^ studies that suggested perturbations in α1 play important roles in accelerating GDP dissociation.

In the third pathway, the interaction between ICL2 of NTR1 and the αN-β1 hinge propagates through the β1 strand and perturbs the GDP phosphate-binding P-loop (α1β1 loop) (Fig. 4d). Such a P-loop perturbation by the αNβ1-ICL2 interaction is also supported by previous structural^3,4,20^ and HDX^37^ studies. This perturbation results in a displacement of the P-loop including the mainchain of residue A41 so that it would no longer be able to make direct interactions with the β-phosphate of GDP (Fig. 4d). To accommodate the displaced P loop, the sidechain of E245 on α2 has rotated by 95° (Fig. 4h). This implies a coupling of P-loop disorder with E245 rotation in the GDP dissociation process, and conversely a role for E245 in maintaining a stable GDP-bound G protein conformation, which coincides with the E245A mutant causing dominant negative effects suggesting a key role of the side chain of E245 in the GDP/GTP exchange process^12,39^. This rotation is not observed in the detergent-embedded NTR1-G_i_ structure, as the P loop adopts a conformation more similar to the one observed in the GDP-G_i_ structure^29^ (Fig. 4h).

Together, this multi-point coordination mechanism leads to complete dissociation of GDP from G_i_ and the creation of a free nucleotide binding pocket for GTP association (Fig. 5).

## Conclusion

Understanding the structural basis for the interaction between GPCRs and G proteins under physiological conditions has been challenging due to the poor stability of the complexes in detergent micelles. Most of the published structures required antibodies/nanobodies and/or engineered G proteins for additional stability, which rendered the complexes incapable of GDP/GTP exchange. Using our recently developed covalently circularized nanodiscs^28^, we have determined two structures, representing different conformational states, of the NT-NTR1-Gα_i1_β_1_γ_1_ complex in a lipid bilayer without the need for external stabilization. These structures reflect the native signaling-competent states of the complex. They show that the sideway movement of TM6, which is considered a signature of active receptors in detergents, is restricted here by the membrane, highlighting the importance of the membrane in modulating the dynamics of GPCR-G protein interactions and the affinity between NTR1 with G_i_. Our structures also allowed us to unravel the interconnected roles of membrane-protein interaction, G-protein activation, and GDP dissociation. The proposed multipartite allosteric mechanism of GDP release reveals a competition between GDP and NTR1 for binding G_i_, which explains why the NTR1-G_i_ complex is stabilized by removal of GDP using apyrase. This observation agrees with a previous NMR study showing that the interaction between NTR1 and Gα is strongest when Gα is nucleotide free^42^. Our study therefore provides new insights into the signal transduction process triggered by GPCR-G protein complex formation and will serve as a model for future studies of GPCR signaling in native lipid bilayers.

## Methods

No statistical methods were used to predetermine sample size. The experiments were not randomized and the investigators were not blinded to allocation during experiments and outcome assessment.

### Preparation of NTR1 in cNDs

Expression and purification of a thermostable variant of rat NTR1 (TM86V-L167R ΔIC3B) was performed as described previously with some modifications^30,43^. This NTR1 variant consists of residues G50-G390, contains a deletion of E273-T290 in intracellular loop (ICL) 3, and has ten stabilizing mutations. Briefly, the full-length fusion protein consisting of maltose-binding protein (MBP), NTR1, and thioredoxin (TrxA) was expressed in Tuner™ (DE3) Competent Cells (Novagen) in LB medium at 37 °C, 200 rpm and induced at an OD_600_ of 0.75 with 1 mM IPTG. Cells were grown for another 24 hours at 20 °C, 160 rpm and harvested by centrifugation (5,000 × *g*, 30 min, 4 °C). Cells were then lysed and solubilized by sonication in buffer containing 100 mM HEPES (pH 8.0), 20% glycerol, 400 mM NaCl, 2.5 mM MgCl_2_, 0.6/0.12% CHAPS/cholesterol, 1.7% n-decyl-β-D-maltopyranoside (DM), 100 mg lysozyme, one tablet of protease inhibitor, and 250 U benzonase. Cell lysate was centrifuged, and the supernatant was mixed with pD-NT resin^43^ pre-equilibrated with 25 mM HEPES (pH 8.0), 10% glycerol, 600 mM NaCl and 0.5% DM at 4 °C for 1 hour. The flow-through from the pD-NT resin was then discarded, and the resin was washed with 25 mM HEPES (pH 7.0), 10% glycerol, 150 mM NaCl, 2 mM DTT and 0.3% DM. The resin was then mixed with 3C protease for 1 hour at 4 °C to cleave off MBP and TrxA from NTR1, as well as NT-NTR1 from pD resin^43^. The resin was washed with 10 mM HEPES (pH 7.0), 10% glycerol, 2 mM DTT and 0.3% DM, which was combined with the flow-through and loaded onto a SP cation exchange chromatography column (GE Healthcare) pre-equilibrated in the same washing buffer. The SP column was washed with 10 mM HEPES (pH 7.7), 10% glycerol, 35 mM NaCl, 2 mM DTT and 1% diheptanoylphosphatidylcholine (DH_7_PC), and then eluted with 10 mM HEPES (pH 7.7), 10% glycerol, 350 mM NaCl, 2 mM DTT and 0.2% DH_7_PC. The eluate was concentrated to below 500 μL and subjected to size-exclusion chromatography on a Superdex 200 10/300 Increase Analytical (S200a) column (GE Healthcare) equilibrated with 10 mM HEPES (pH 7.7), 150 mM NaCl, 2 mM DTT, 0.1% DH_7_PC and 0.1 μM NT. Fractions containing NTR1 were collected and mixed with a 3:2 molar ratio of 1-Palmitoyl-2-oleoyl-phosphatidylcholine (POPC) to 1-palmitoyl-2-oleoyl-phosphatidylglycerol (POPG) solubilized in 100 mM sodium cholate at a NTR1:lipid molar ratio of 1:160. The mixture was incubated on ice for 30 min before addition of cNW9 at a cNW9:NTR1 molar ratio of 4:1 followed by another 30 min incubation on ice. The mixture was then treated with 5% volume of Bio-Bead SM-2 resin (Bio-Rad) with shaking on ice for 15 min, followed by addition of another 20% volume of Bio-Beads every 20 min for detergent removal. After two-hour incubation with Bio-Beads, the flow-through was then subjected to size-exclusion chromatography with a S200a column equilibrated in 20 mM sodium phosphate (pH 6.9), 50 mM NaCl, 1 mM DTT, 0.5 mM EDTA, 0.1 μM NT. Fractions containing NTR1-cND were concentrated to below 500 μL and incubated at 42 °C for 24 hours, followed by filtration through 0.22 μm cut-off filters. The flow-through was subjected to another round of size-exclusion chromatography. Fractions were pooled, concentrated and stored at 4°C.

### Preparation of Gα_i1_β_1_γ_1_ in micelles and cNDs

G protein composed of Gα_i1_, Gβ_1_ and Gγ_1_ was expressed and purified as detailed before^30,44^. Briefly, *Spodoptera frugiperda* (Sf9) were grown in suspension in ESF921 medium (Expression Systems, California), infected at a density of 2-3 × 10^6^/mL with a single baculovirus encoding all three subunits (Gα_i1_β_1_γ_1_), harvested within 72 hours post inoculation, and stored at -80 °C until use.

Cells were lysed in 10 mM HEPES (pH 7.4), 20 mM KCl, 10 mM MgCl_2_, 10 μM GDP, 2 mM β-mercaptoethanol (β-ME), and 1 protease inhibitor tablet with sonication. The suspension was then ultra-centrifuged at 180,000 × *g* for 45 min at 4 °C. The membrane pellet was solubilized in 50 mM HEPES (pH 7.4), 150 mM NaCl, 10 mM MgCl_2_, 10 μM GDP, 2 mM β-ME, 10% glycerol, 1 protease inhibitor tablet, 1.2% DM at 4 °C for 3 hours. The suspension was ultra-centrifuged again and the supernatant was purified through Ni-NTA resin^20^. The eluate was concentrated and run through a Superdex 200 prep 16/60 column (S200p; GE Healthcare) equilibrated in 20 mM HEPES (pH 7.4), 100 mM NaCl, 0.1 mM MgCl_2_, 4 mM β-ME, and 0.5% DM. Fractions containing Gα_i1_β_1_γ_1_ were pooled and concentrated to 10 mg/mL, flash-frozen in liquid nitrogen and stored at -80 °C.

Gα_i1_β_1_γ_1_-cNDs were prepared similarly as for NTR1-cNDs. After Bio-Bead removal, the Gα_i1_β_1_γ_1-_ -cNDs were purified through Ni-NTA to remove empty cNDs, followed by S200a chromatography to remove aggregates. Fractions containing pure Gα_i1_β_1_γ_1_-cNDs were collected, concentrated, and stored at 4°C.

### Complex formation of NT-NTR1-Gα_i1_β_1_γ_1_ in cNDs

Purified Gα_i1_β_1_γ_1_ in micelle was diluted in buffer A (20 mM HEPES (pH 6.9), 50 mM NaCl, 5 mM CaCl_2_, 1 mM DTT, 0.1 μM NT) until the DM concentration dropped below 0.08% (the critical micelle concentration of DM), and mixed with NTR1-cND at 1:1 molar ratio. The mixture was incubated on ice for 30 min, followed by addition of Bio-Beads at 10% volume every 30 min. The mixture was incubated on ice with shaking for a total of 2 hours and then the Bio-Beads were removed. Apyrase, diluted with buffer A and pretreated with Bio-Beads for 30 min on ice, was added to the complex at 1 U/mL concentration. The mixture was incubated at 4 °C overnight, and then subjected to a S200a SEC column equilibrated in 20 mM sodium phosphate (pH 6.9), 50 mM NaCl, 1 mM DTT, 0.5 mM EDTA, 0.1 μM NT. Peak fractions were characterized with SDS-PAGE and negative-stain EM. The fractions containing NT-NTR1-Gα_i1_β_1_γ_1_ in cNDs were used for cryo-EM structure determination.

### Circular Dichroism (CD) spectroscopy

CD spectra were measured on a JASCO J-815 spectropolarimeter equipped with a Peltier cell temperature controller. Both spectrum scan measurement and variable temperature measurement were carried out for the following samples: NTR1 in micelles, NTR1-cNDs, Gα_i1_β_1_γ_1_ in micelles, Gα_i1_β_1_γ_1_-cNDs, and NTR1-Gα_i1_β_1_γ_1_ in cNDs. Spectrum scan measurements were performed at 20 °C, before and after variable temperature measurements, collecting data from 260 nm to 190 nm. Variable temperature measurements were carried out at 220 nm increasing temperature from 20 °C to 95 °C at a rate of 1 °C/min. *Spectrum Manager 2* software was used to analyze the transition temperature for each sample.

### Binding affinity and kinetics measurement

Binding affinity and kinetics between NTR1 and Gα_i1_β_1_γ_1_ in detergent micelles and cNDs were measured using MicroScale Thermophoresis (MST) and Biolayer Interferometry (BLI).

For MST, the measurements were performed on a Monolith NT.115 system (NanoTemper Technologies). We measured the fluorescence signal from Gα_i1_β_1_γ_1_ by using the Monolith His-Tag Labeling Kit RED-tris-NTA 2nd Generation kit (NanoTemper Technologies). The samples were prepared in a buffer containing 20 mM sodium phosphate (pH 6.9), 50 mM NaCl, 0.05% DH_7_PC for cND titrations and 0.2% DH_7_PC for titrations in detergent micelle. The concentration of DH_7_PC for cND titrations is below its critical micellar concentration. The experiments were carried out as fast as possible (within 1-2 minute for sample preparation) to prevent degradation of Gα_i1_β_1_γ_1_. The concentration of Gα_i1_β_1_γ_1_ was constant at 10 nM. NT-NTR1 in DH_7_PC, NT-NTR1-cND, or empty cND was titrated in two-fold dilution steps beginning at 4 µM. For the measurement the samples were filled into premium-coated capillaries. The measurement was performed at 2 % LED and 20 % MST power, 30 sec Laser-On, and 5 sec Laser-Off. Fluorescence was excited at 605–645 nm, and emission was detected at 680–685 nm. The results were analyzed using the MO Affinity Analysis software (NanoTemper Technologies). The dissociation constant (K_D_) was then determined using a single-site model for data fitting.

BLI experiments were performed on an Octet RED384 (ForteBio, California) using Anti-His antibody-coated Dip and Read Biosensors (HIS1, ForteBio) and 384 well plates (ForteBio) with 60 µL volume. 500 nM of His-tagged Gα_i1_β_1_γ_1_ was bound for 5 min in a binding buffer consisting of 20 mM HEPES (pH 7.4), 100 mM NaCl, 0.1 mM MgCl_2_, 4 mM β-ME, and 0.5% DM. To test for nonspecific binding of His-tagged Gα_i1_β_1_γ_1_, reference tips were incubated in buffer alone. The tips were washed with buffer for 2 min to obtain a baseline reading and then transferred to wells in various concentrations of NT-NTR1-cND (4, 2, 1, 0.5, 0.25 µM) in buffer containing 20 mM sodium phosphate (pH 6.9), 50 mM NaCl, 1 mM DTT, 0.5 mM EDTA, 0.1 μM NT for 5 min. After measuring the association phase, tips were moved to wells containing buffer with and without GTPγS, and dissociation was measured for 5 min. The data were processed and analyzed using the Octet data analysis software version 11.0 (ForteBio). Association-dissociation curves for each concentration were fit to a 2:1 model.

### Nuclear Magnetic Resonance (NMR) spectroscopy

Uniformly ^15^N-labeled NT-NTR1 in POPC/POPG (3:2) cNW9 nanodiscs at 200 μM alone and in complex with unlabeled Gα_i1_β_1_γ_1_ at a molar ratio of 5:1 were prepared as described above in NMR buffer (20 mM sodium phosphate (pH 6.9), 50 mM NaCl, 1 mM DTT, 0.5 mM EDTA, 10% D_2_O). Two-dimensional Transverse Relaxation Optimized Spectroscopy (TROSY) Heteronuclear Single Quantum Coherence (HSQC) were collected with 2000 scans, 200 increments at 45 °C on a Bruker 800-MHz spectrometer equipped with a TXO cryogenic probe. TROSY HSQC measurements were repeated for NT-NTR1-cND on an Agilent 700-MHz spectrometer to verify that NT-NTR1-cND stays intact after long data acquisition in the magnet at 45 °C. Data were processed using the NMRPipe software package^45^.

### Functional Assay

Ligand-induced IP1 (a metabolite of IP3) accumulation was measured in transiently transfected HEK293T/17 cells as described before^46^. Wild type rNTR1 or mutants thereof were directly subcloned into a mammalian expression vector containing an N-terminal SNAP-tag (pMC08). 24 hrs after transfection, cells were washed with PBS, detached with Trypsin-EDTA (Sigma) and resuspended in assay buffer (10 mM HEPES pH 7.4, 1 mM CaCl_2_, 0.5 mM MgCl_2_, 4.2 mM KCl, 146 mM NaCl, 50 mM LiCl, 5.5 mM glucose, 0.1% (w/v) BSA). Cells were seeded at 20,000 cells per well in white 384-well plates (Greiner) and incubated for 2 hrs at 37 °C with a concentration range of NT_8–13_ (Anawa) diluted in assay buffer. IP1 accumulation was measured using the HTRF IP-One kit (Cisbio) according to the manufacturer’s protocol. To confirm cell surface expression of NTR1 and its mutants, transfected cells were plated on poly-D-lysine treated 384-well plates (Greiner) at 20,000 cells/well in growth medium. The following day, medium was removed and cells were incubated with 50 nM SNAP-Lumi4-Tb (CisBio) in labelling buffer (CisBio) for 2 hrs at 37 °C. Thereafter, cells were washed 4 times with wash buffer (20 mM HEPES pH 7.5, 100 mM NaCl, 3 mM MgCl_2_ and 0.2% (w/v) nonfat milk). Fluorescence intensity of Tb^3+^-labelled receptors was measured on an Infinite M1000 fluorescence plate reader (Tecan) with an excitation wavelength of 340 nm and emission wavelength of 620 nm. To generate concentration-response curves, data were normalized to receptor expression at the cell surface and to response of NTR1 at maximal ligand concentration and were analysed by a non-linear curve fit in GraphPad Prism.

### Negative-stain microscopy

3 µL of NT-NTR1-Gα_i1_β_1_γ_1_-cND complex at a concentration of 0.02 mg/mL was applied onto a glow-discharged continuous carbon grid (Electron Microscopy Sciences, Inc.). After two minutes of adsorption, the grid was blotted with filter paper to remove the excess sample, immediately washed twice with 50 µL of MiliQ water, once with 50 µL 0.75% uranyl formate solution and incubated with 50 µL of 0.75% uranyl formate solution for an additional one minute. The grid was then further blotted with filter paper to remove the uranyl formate solution, air-dried at room temperature, and examined with a Tecnai T12 electron microscope (Thermo Fisher Scientific) equipped with an LaB6 filament and operated at 120-kV acceleration voltage, using a nominal magnification of 52,000× at a pixel size of 2.13 Å.

### Cryo-EM sample preparation

Cryo-EM grids were prepared using a Vitrobot Mark IV (Thermo Fisher Scientific). 3 μL of NT-NTR1-Gα_i1_β_1_γ_1_-cND at a concentration between 1.5 mg/mL to 1.7 mg/mL was applied onto glow discharged C-flat holy carbon grids (R1.2/1.3, 400 mesh copper, Electron Microscopy Sciences) or Quantifoil holy carbon grids (R1.2/1.3, 400 mesh gold, Quantifoil Micro Tools). The grids were blotted for 7.5 s with a blot force of 16 and 100% humidity before being plunged into liquid ethane cooled by liquid nitrogen.

### Cryo-EM data collection

Images of NT-NTR1-Gα_i1_β_1_γ_1_-cND were acquired on Titan Krios I at the Harvard Cryo-EM Center for Structural Biology equipped with a BioQuantum K3 Imaging Filter (slit width 20 eV) and a K3 direct electron detector (Gatan) and operating at an acceleration voltage of 300 kV. Images were recorded at a defocus range of -1.2 µm to -2.5 µm with a nominal magnification of 105,000×, resulting in a pixel size of 0.825 Å. Each image was dose-fractionated into 38 movie frames with a total exposure time of 1.5 s, resulting in a total dose of ∼57 electrons per Å^2^. SerialEM was used for data collection^47^.

### Image processing

A total of 23,677 movie stacks, which were collected during two sessions, were motion corrected and electron-dose weighted using MotionCor2 (ref.^48^). Parameters of the contrast transfer function were estimated from the motion-corrected micrographs using CTFFIND4 (ref.^49^). To generate a reference, particles from 10 micrographs were picked manually in EMAN2.2 (ref.^50^), crYOLO^51^ was then trained for picking particles automatically. All subsequent 2D and 3D analyses were performed using RELION-3.0 or RELION-3.1-beta^52^.

1,726,457 particles were selected after several rounds of 2D classification from 4,367,542 auto-picked particles. Density map of the human NTR1 in complex with the agonist JMV449 and the heterotrimeric G_i1_ protein (EMDB-20180)^8^ was low-pass filtered to 20 Å and used as the initial model for the first round of 3D classification, yielding five different classes. Two classes of the NT-NTR1-Gα_i1_β_1_γ_1_-cND complex were relatively better resolved and particles from these two classes were subject to 3D refinements. Bayesian polishing was then performed, followed by 3D refinement and post-processing, yielding two density maps at resolutions of 4.3 Å (canonical state) and 4.5 Å (noncanonical state), respectively. To further improve the resolution of the core of the complex, masks excluding the nanodisc and the AHD were applied during the 3D refinement, yielding the 4.1 Å (canonical state) and 4.2 Å (noncanonical state) density maps, respectively. Per-particle CTF refinement was performed but did not lead to an improvement in map resolution or quality.

### Model building and refinement

The crystal structures of NT-NTR1 complex (PDB: 4BUO)^30^ and G protein heterotrimer Gα_i1_β_1_γ_2_ (PDB: 1GP2)^29^ and the cryo-EM structure of Gα_T_β_1_γ_1_ (PDB: 6OY9)^9^ were fitted into the density map of the canonical NT-NTR1-Gα_i1_β_1_γ_1_-cND complex using the Fit in Map function of Chimera^53^. The α_i1_β_1_ subunits of Gα_i1_β_1_γ_2_ and γ_1_ subunit of Gα_T_β_1_γ_1_ were merged with the NT-NTR1 structure and the amino acids were modified in Coot version 0.9-pre to match our constructs^54^. The amino acids F291-R299 of NTR1 of the canonical state were mutated to poly-alanine due to the lack of sidechain densities. The model was manually adjusted and refined in Coot with torsion, planar peptide, trans peptide and Ramachandran restraints applied. For the noncanonical state, the subunits of the refined atomic model of the canonical state were fitted into the density map as separate rigid bodies. The model was manually adjusted and refined in Coot. For both states, the AHD was extracted from the crystal structure of the human Gα_i1_ (PDB: 3UMR) and docked into the density as a rigid body using Chimera.

Models were refined with Phenix.real_space_refine^55^. The AHD was not refined due to the lack of sidechain information for this domain. During refinement, the resolution limit was set to match the map resolution determined by the FSC=0.143 criterion in post-processing. Secondary structure, Ramachandran, rotamer, and reference restraints from the JMV449-NTR1-G_i_-scFv16 complex (PDB 6OS9)^8^ were applied throughout refinement. The final models were validated using MolProbity v.4.3.1 (ref.^56^) with model statistics provided in Table S1.

### Molecular dynamics simulations

The molecular system for the molecular dynamics (MD) simulations was prepared based on the canonical state structure of NT-NTR1-Gα_i1_β_1_γ_1_-cND which was preprocessed with Maestro from Schrödinger^57,58^. Bond orders were assigned, hydrogens added, disulfide bonds created, and het states generated at pH 7.0±2.0. The sidechains of residues 291 to 299 were assigned and the truncated residues 273 to 290 in NTR1 construct were added with the Crosslink Proteins tool of Maestro^57,58^.

The membrane and solvent environment, as well as the input files for Amber were generated using the Membrane Builder tool of CHARMM-GUI^59,60^. The terminal groups of each chain were patched with standard N-terminus and C-terminus patch residues, except for the N-terminus of Gα for which a GLYP patch residue was used. For orienting the complex appropriately, the PPM (Positioning of Proteins in Membrane) server of the OPM (Orientations of Proteins in Membranes) database was used^61^. A lipid bilayer containing a total of 527 lipids, composed of a 3:2 molar ratio of POPC to POPG, was added to the aligned complex with Membrane Builder^59,60^. A rectangular solvation box was added by adding water layers of at least 22.5 Å above and below the membrane. The system was ionized and neutralized by adding 50 mM of sodium and chloride ions. The resulting system contained a total of 286,109 atoms.

In total, 12 simulations of the prepared system were run using Amber18 (ref.^62^). The Amber FF14SB^63^ and Amber Lipid17 (ref.^64^) force fields were used for the proteins and the lipid bilayer, respectively. The TIP3P model^65^ was used for the water molecules. During the energy minimization, 2500 steps of steepest descent followed by 2500 steps of conjugate gradient were carried out. The equilibration steps were carried out according to the standard Membrane Builder protocols^66^. The production MD simulations were carried out at 310 K and 1 bar in an NPT ensemble using a Monte Carlo barostat and a Langevin thermostat. The cutoff for the nonbonded interactions was set to 10 Å, and the particle mesh Ewald method was used for the long-range electrostatic interactions. Hydrogen mass repartitioning was enabled, and a time step of 4 fs applied. Postprocessing was carried out with AmberTools 18 and VMD 1.9.4 (ref.^62^) The simulation lengths of the runs were between 600 ns and 1 µs.

## Acknowledgments

Cryo-EM data were collected at the Harvard Cryo-Electron Microscopy Center for Structural Biology. We thank F. Koh, P. Egloff, P. Heine, M. Hillenbrand, and J. Schöppe for their contribution to the early stages of this project, S. Sterling, R. Walsh, and Z. Li for microscopy support, SBGrid for computing support, M. Deluigi for supervising the signaling experiments, and R. Walker, K. Bayer, P. Imhof and M. Bagherpoor for their advice and discussions regarding the molecular dynamics simulations. M.G. is supported by a Merck-BCMP fellowship. A.B. is supported by the International Retinal Research Foundation, the E. Matilda Ziegler Foundation for the Blind, the Richard and Susan Smith Family Foundation, and the Pew Charitable Trusts. We acknowledge support by NIH grants GM129026 and AI037581 to G.W. and GM131401 to M.L.N. G.W. and A.P. are supported by HFSP RGP0060/2016. A.P. is supported by Swiss National Science Foundation grant 31003A_182334.

## Author contributions

M.Z. developed the protocol for making NT-NTR1-Gα_i1_β_1_γ_1_-cND complexes, prepared samples, collected negative-stain EM images, and performed biophysical experiments. M.G. prepared cryo-EM grids, obtained and processed the data, and built and refined the atomic models. M.Z. and Z.W. performed binding experiments. C.G. performed MD simulations. J.Y. expressed Gα_i1_β_1_γ_1_. H.W. obtained and processed cryo-EM data. M.Z. and J.S. performed NMR experiments. C. K., L. Me and L. Mo made constructs and performed signaling experiments. A.P. designed and supervised the signaling experiments. F.H. and G.W. initiated the project. A.B., M.L.N., and G.W. designed and supervised the project. M.Z. wrote the paper. M.Z., M.G., Z.W., C.G., J.Y., C. K., A.P., A.B., M.L.N., and G.W. edited the paper.

## Data and materials availability

Structural data have been deposited into the Worldwide Protein Data Bank (wwPDB) and the Electron Microscopy Data Bank (EMDB). The EM density maps for the canonical and noncanonical states of NT-NTR1-G_i_ complex in lipid nanodiscs have been deposited under accession codes EMD-XXXX and EMD-XXXX. The masked EM density maps excluding the nanodisc and AHD of the canonical and noncanonical states have been deposited under accession codes EMD-XXXX and EMD-XXXX. The corresponding atomic models have been deposited under accession codes YYYY and YYYY. Other data are available upon reasonable request.

---

**Extended Data Fig. 1.**
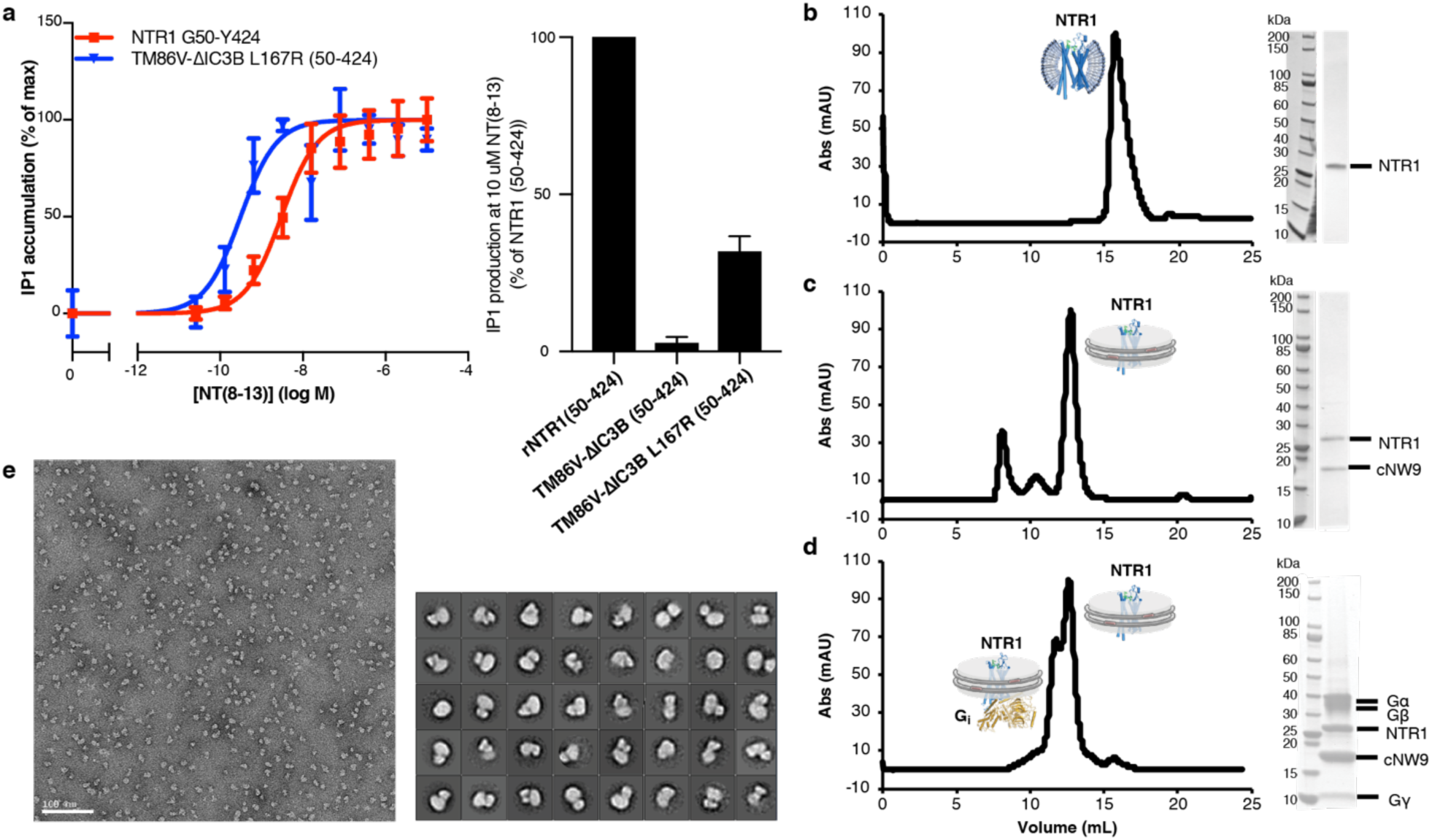
Signaling competency and preparation of NT-NTR1-G_i_ complex in cNDs. **a**, Signaling competency of NTR1 constructs. Wild-type NTR1 (50-424) or NTR1 variants were transiently transfected into HEK293T/17 cells, and activation of Gα_q_ signaling was quantified by measuring of inositol-1-phosphate (IP1) accumulation after stimulation with NT_8–13_. Data were normalized to receptor expression at the cell surface and represented as mean ± SEM of 4 independent experiments performed in duplicate. Left, dose dependent IP1 production expressed as percentage of IP1 accumulation at maximal ligand concentration. Fitting of the curves result in EC50 of 2.664 nM for wild-type NTR1 and 0.2188 nM for TM86V ΔIC3B L167R. Right, bar graph showing IP1 production level at 10 μM agonist NT_8-13_. The NTR1 variant TM86V ΔIC3B lacking the L167R back mutation exhibits no IP1 production, suggesting the critical role of R167^3.50^ in signal transduction. **b-d**, Size-exclusion chromatograms and corresponding SDS-PAGE gels for NTR1 in DH_7_PC detergent micelles (**b**), NTR1 in POPC/POPG cNW9 nanodiscs (**c**) and NTR1-G_i_ complex in POPC/POPG cNW9 nanodiscs (**d**). **e**, Fractions corresponding to the NT-NTR1-G_i_ complex in (**d**) were analyzed by negative-stain EM, and then used for cryo-EM structure determination. Top, representative negative-stain EM micrograph of NT-NTR1-G_i_ complexes in cNDs. Bottom, 2D class averages.

**Extended Data Fig. 2.**
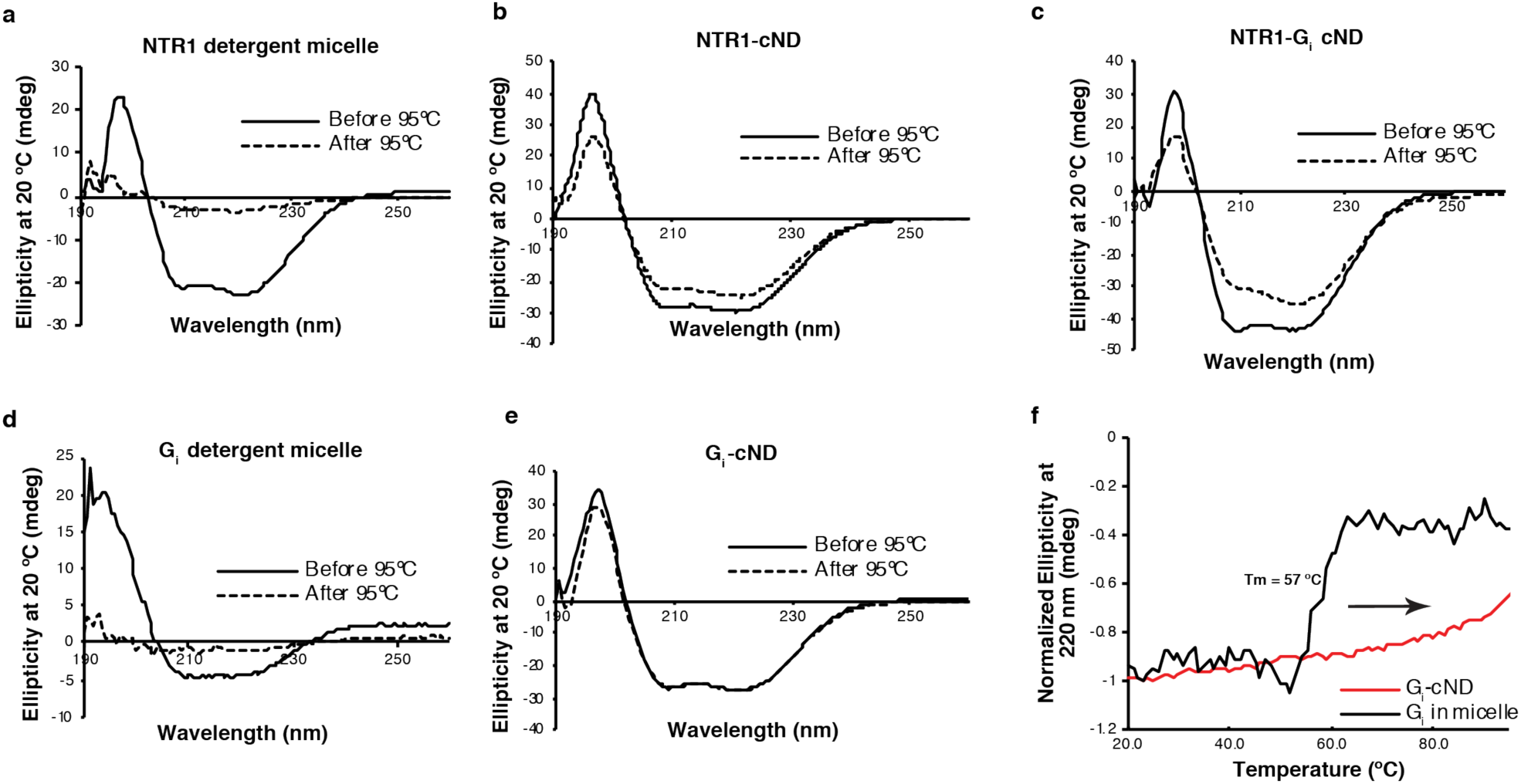
Thermostability enhancement of NTR1, G_i_, and NTR1-G_i_ complexes by incorporation into cNDs. **a-e**, Circular Dichroism (CD) spectra at 20 °C before (solid line) and after (dashed line) treatment at 95 °C of (**a**) NTR1 in DH_7_PC detergent micelles; (**b**) NTR1 in cNDs; (**c**) NTR1-G_i_ complex in cNDs; (**d**) G_i_ in detergent micelle; (**e**) G_i_ in cNDs. G_i_ was reconstituted into cNDs by incubation with POPC/POPG lipid, cNW9, and cholate, followed by detergent removal and size-exclusion chromatography. **f**, Temperature-dependent CD signals of G_i_ in detergent micelles (black) and cNDs (red) at 220 nm. The melting temperature (Tm) of cNDs is 93 °C (data not shown) and therefore does not affect transitions before this temperature. NTR1 and G_i_ account for at least 50% of CD signals even in the presence of cNDs. NTR1 in detergent micelles irreversibly unfolds during temperature increase with a Tm of 62 °C. In contrast, NTR1-cND changes structure around 80 °C and does not lose much secondary structure after decreasing temperature to 20 °C. Similar observations were made for G_i_, where the protein irreversibly and completely unfolds with Tm of 57 °C in detergent micelles but displays no clear transition temperature in cNDs even until 95 °C. For the NTR1-G_i_ complex in cND, only mild unfolding was observed around 82 °C. These observations indicate that lipid bilayers improve the stability of NTR1, G_i_ and NTR1-G_i_ complexes relative to detergent micelles.

**Extended Data Fig. 3.**
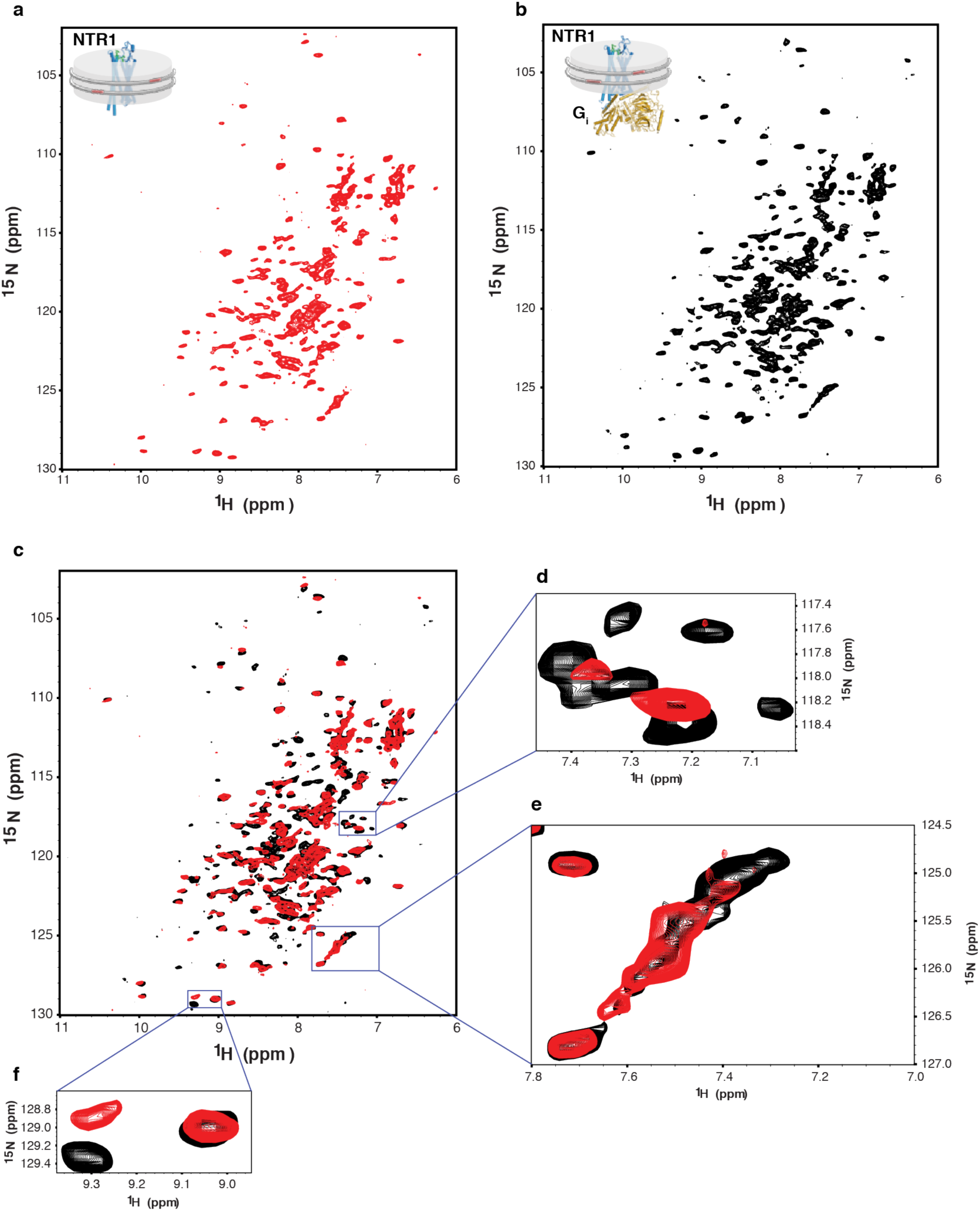
Characterization of the interaction between NT-NTR1 and G_i_ in cNDs by two-dimensional ^1^H, ^15^N-TROSY HSQC NMR spectroscopy. **a-b**, NMR spectrum of ^15^N-labeled NT-NTR1 in cNDs in the absence (**a**) and presence (**b**) of G_i_. **c**, Overlay of (**a**) (red) onto (**b**) (black) showing structural and dynamical changes of NT-NTR1 upon binding to G_i_ in cNDs. **d**, A region showing conformational stabilization of NTR1. More peaks could be observed in the presence of G_i_, suggesting that NTR1 is highly dynamic in the absence of G_i_ and resonances are averaged out among a wide range of conformers resulting in low signal-to-noise ratio and even disappeared peaks. Upon interaction with G_i_, NTR1 is stabilized into fewer conformers and becomes less dynamic, which leads to better signal-to-noise ratio and more resonances being observable. **e**, A region showing dynamically slow-exchange shift of NTR1 upon interaction with G_i_. **f**, A region showing chemical shift perturbation of NTR1, suggesting conformational change of NTR1 upon binding to G_i_ in cNDs.

**Extended Data Fig. 4.**
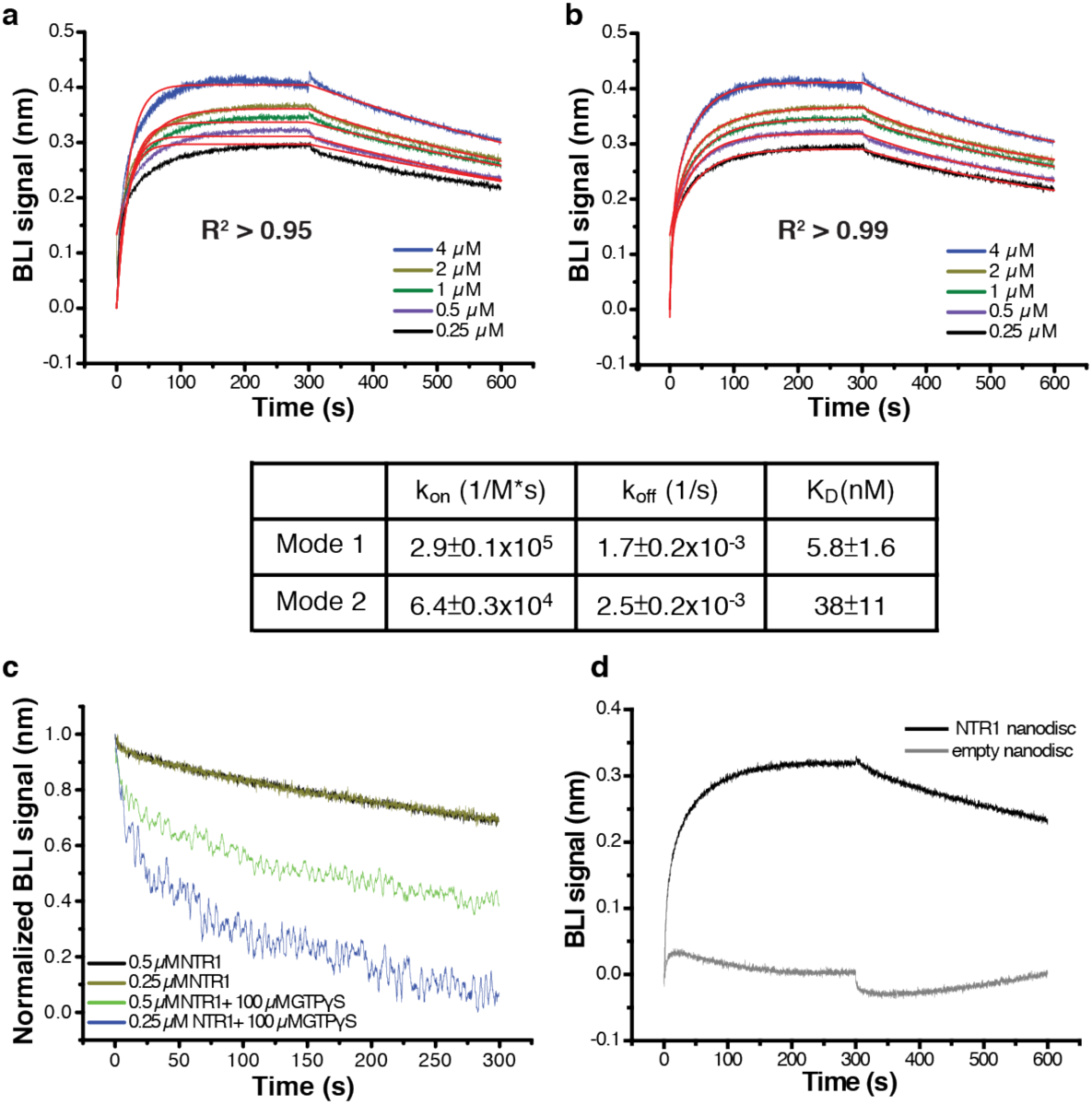
Characterization of the binding kinetics between NT-NTR1 and G_i_ in cNDs. **a-b**, Fitting of Bio-Layer Interferometry (BLI) traces of G_i_ binding to NT-NTR1-cND using one binding mode (**a**) and two binding mode (**b**) shows better fitting using two binding mode. Below, a table showing k_on_, k_off_ and K_D_ from the two binding mode fitting. **c**, Dissociation between G_i_ and NT-NTR1-cND in the absence (black and brown) and presence (green and blue) of GTPγS, showing faster dissociation of the complex in the presence of GTPγS, suggesting the capability of the NTR1-Gα_i1_β_1_γ_1_ complex in cNDs to perform GDP/GTP exchange. **d**, Association and dissociation kinetics of G_i_ binding to NT-NTR1-cND (dark) and empty cND (gray), showing much slower association and faster dissociation of G_i_ binding to empty cND compared to NT-NTR1-cND, suggesting that interaction between G_i_ and NT-NTR1-cND is driven by G_i_ binding to NTR1 rather than to the nanodisc.

**Extended Data Fig. 5.**
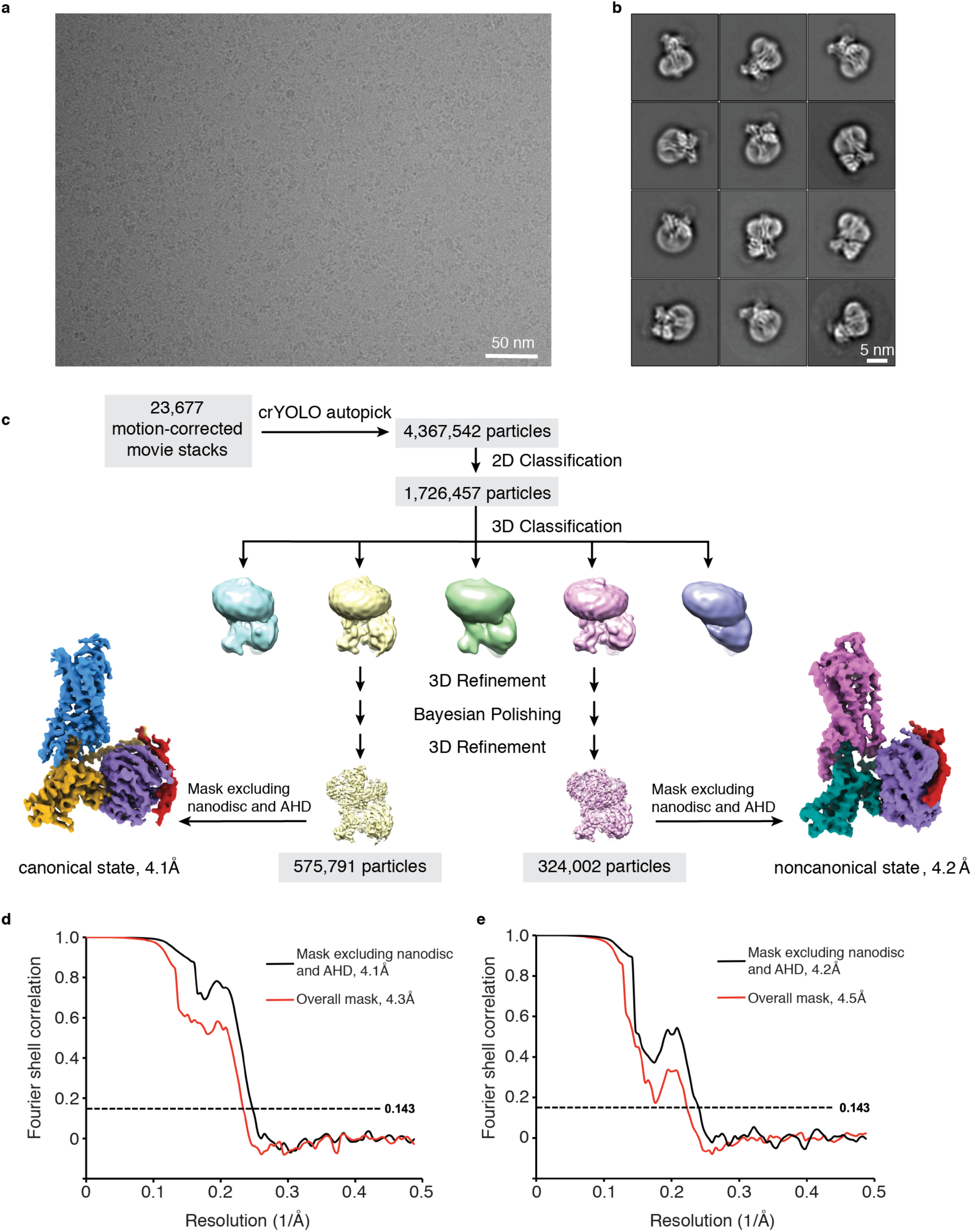
Cryo-EM data processing. **a**, Representative micrograph showing the distribution of NT-NTR1-G_i_-cND particles in vitreous ice. **b**, Selected two-dimensional class averages showing secondary structure features. The cND has an approximate diameter of 9 nm. **c**, Simplified flow chart of the cryo-EM processing. Two datasets were collected and processed similarly; the number of particles shown here are a conflation of both datasets. Two well-resolved classes corresponding to canonical and noncanonical states were identified. Further rounds of classification did not improve the resolution or map quality. **d-e**, Fourier shell correlation (FSC) curves for the canonical state (**d**) and noncanonical state (**e**) with masks that either include or exclude the cND and AHD.

**Extended Data Fig. 6.**
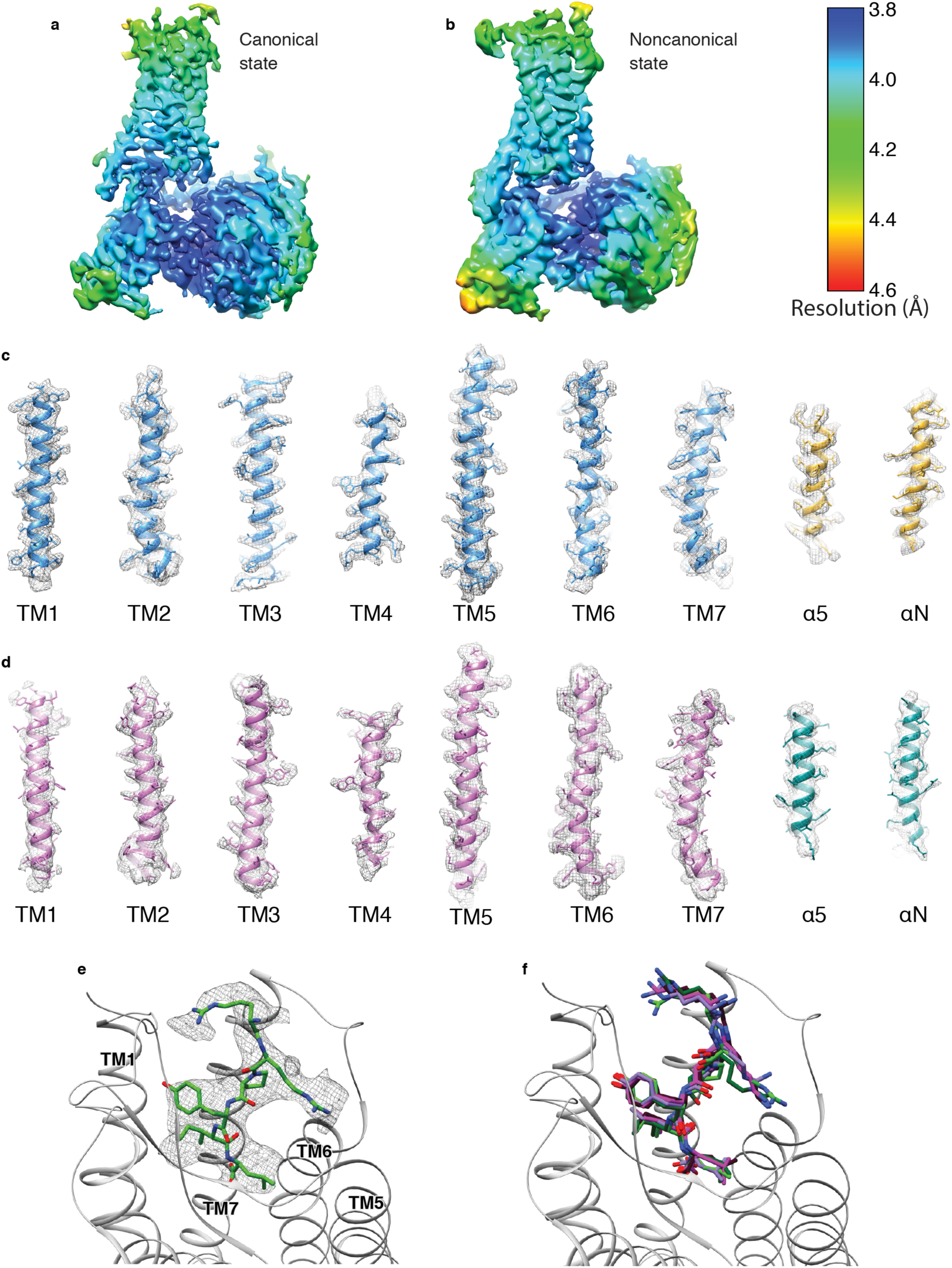
Cryo-EM density. **a-b**, Local resolution of the NT-NTR1-G_i_ complex in canonical state (**a**) and noncanonical state (**b**). The local resolution is calculated in Relion. **c-d**, Density and model for the transmembrane helices of NTR1 and the α5 and αN helices of Gα_i1_ in the canonical state (**c**) and noncanonical state (**d**). **e**, Density and model for NT_8-13_. **f**, Superposition of the atomic models of NT_8-13_ from the NT-NTR1-G_i_-cND complex in the canonical (light green), and noncanonical state (dark green) with NT from the NT-NTR1 crystal structure (purple; PDB 4XEE) and JMV449 (a NT analog) from the NTR1-G_i_-micelle complex in the canonical (magenta; PDB 6OS9) and noncanonical state (dark red; PDB 6OSA).

**Extended Data Fig. 7.**
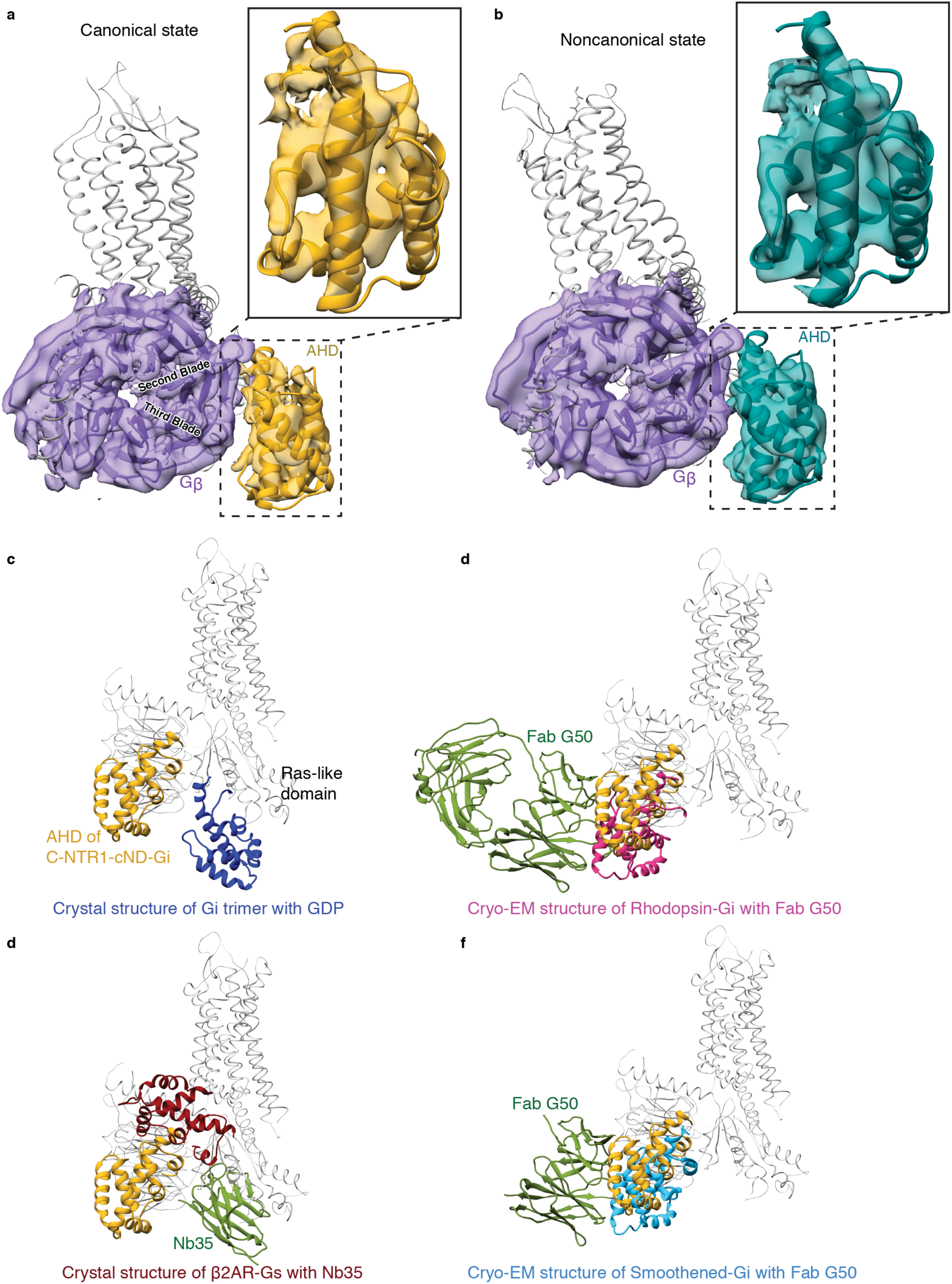
Structure of the α-helical domain (AHD). **a**, Density maps and models showing the interaction between Gβ_1_ (purple) and Gα_i1_ AHD (gold) in the canonical state. Zoom-in view of the Gα_i1_ AHD is shown. **b**, Density maps and models showing the interaction between Gβ_1_ (purple) and Gα_i1_ AHD (dark green) in the noncanonical state. Zoom-in view of the Gα_i1_ AHD is shown. The models in (**a**) and (**b**) are superposed on the Gβ_1_ subunits and are shown in the same view. AHD in both states interacts with the second and third blades of Gβ_1_. **c-f**, Comparison of the AHD of the canonical state NT-NTR1-Gi-cND (gold) with **c**, A crystal structure of GDP-G_i_ (blue; PDB 1GP2), **d**, A crystal structure of β_2_AR-G_s_ with nanobody Nb35 (AHD is dark red and Nb35 is green; PDB 3SN6), **e**, A cryo-EM structure of Rhodopsin-G_i_ with Fab G50 (AHD is pink and Fab G50 is green; PDB 6CMO), and **f**, A cryo-EM structure of Smoothened-G_i_ with Fab G50 (AHD is light blue and Fab G50 is green; PDB 6OT0). The models are superposed on the Gα Ras-like domain.

**Extended Data Fig. 8.**
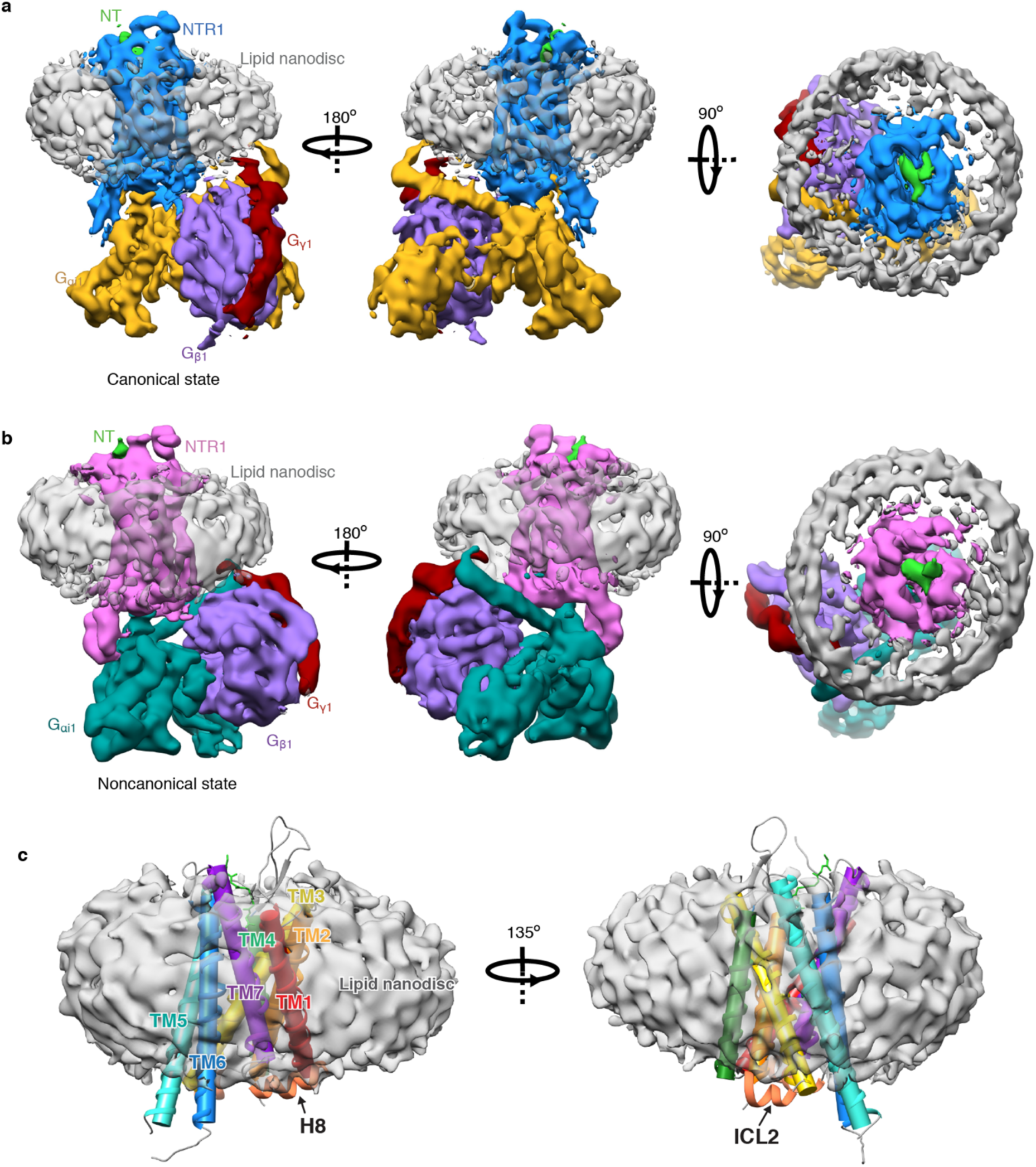
Cryo-EM structure of the NT-NTR1-G_i_ complex in lipid nanodiscs and the interaction with lipid. **a**, Three views of the cryo-EM density map of the NT-NTR1-G_i_-cND complex in the canonical state. **b**, Three views of the cryo-EM density map of the NT-NTR1-G_i_-cND complex in the noncanonical state. The maps in panels (**a**) and (**b**) are low-pass filtered to 5 Å and colored by subunit. **c**, Two views of NT-NTR1 surrounded by nanodisc density. The transmembrane helices are shown in cylinder representation using the rainbow coloring scheme. ICL2 and helix H8 are partially submerged in lipid.

**Extended Data Fig. 9.**
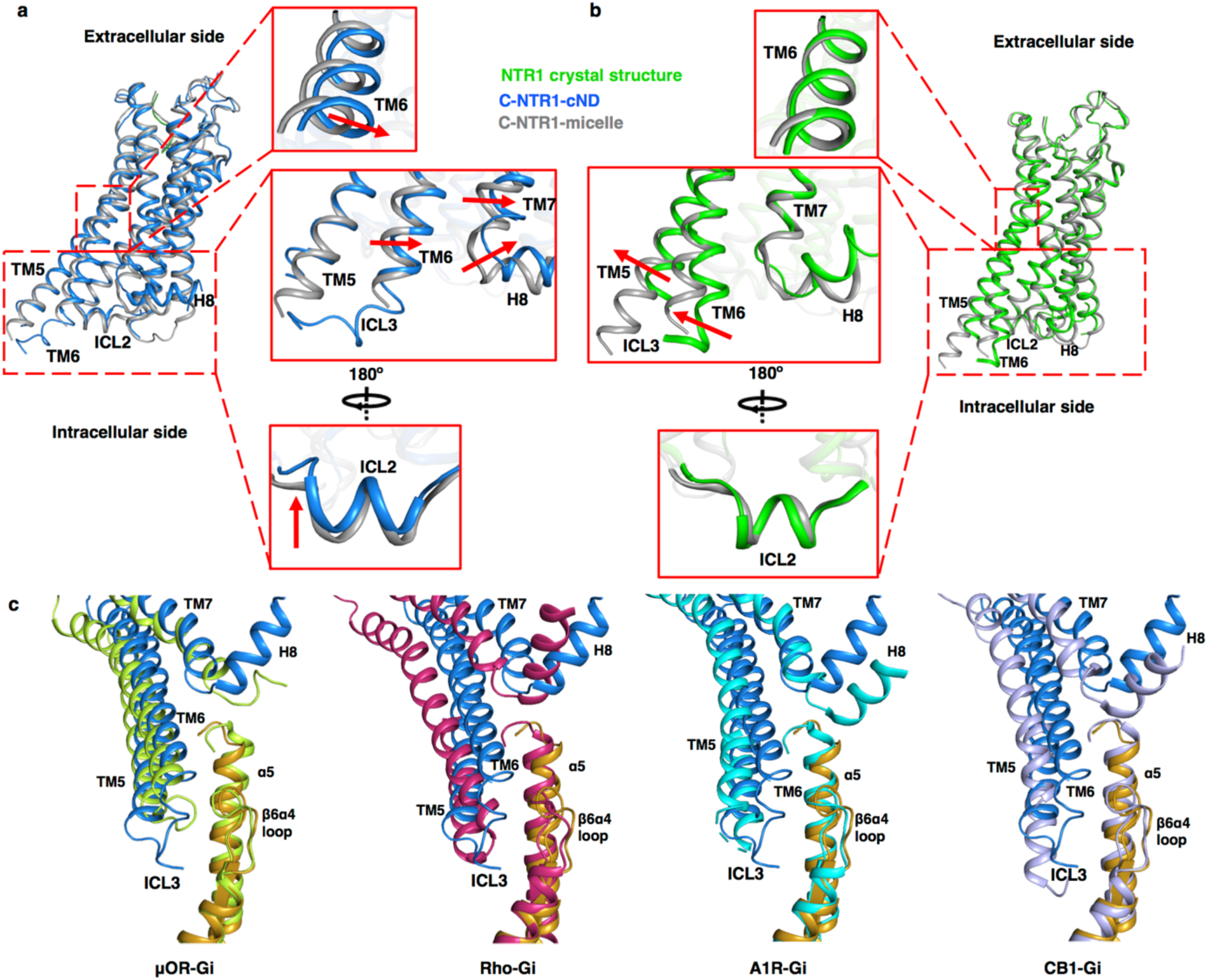
Impact of lipid bilayer on the structure of NTR1. **a**, Comparison between the cryo-EM structures of the canonical states of NTR1 (with G_i_) in lipid bilayer (blue) and detergent (gray). **b**, Structural comparison between the crystal structure of NTR1 in detergent (green) and the cryo-EM structure of the canonical state of NTR1 in complex with G_i_ in detergent (gray). The atomic models in (**a**) and (**b**) are superposed on NTR1. **c**, Comparison of the localization of TM5-TM6 relative to α5 helix of Gα in class A GPCR-G_i_ complex structures, including the canonical state NTR1 (blue) in complex with G_i_ (gold) structure reported in the current study, μOR-G_i_ (lime green; PDB 6DDE), Rho-G_i_ (hot pink; PDB 6CMO), A_1_R-G_i_ (cyan; PDB 6D9H), and CB1-G_i_ (purple; PDB 6N4B). The models are superposed on the Ras-like domain of Gα.

**Extended Data Fig. 10.**
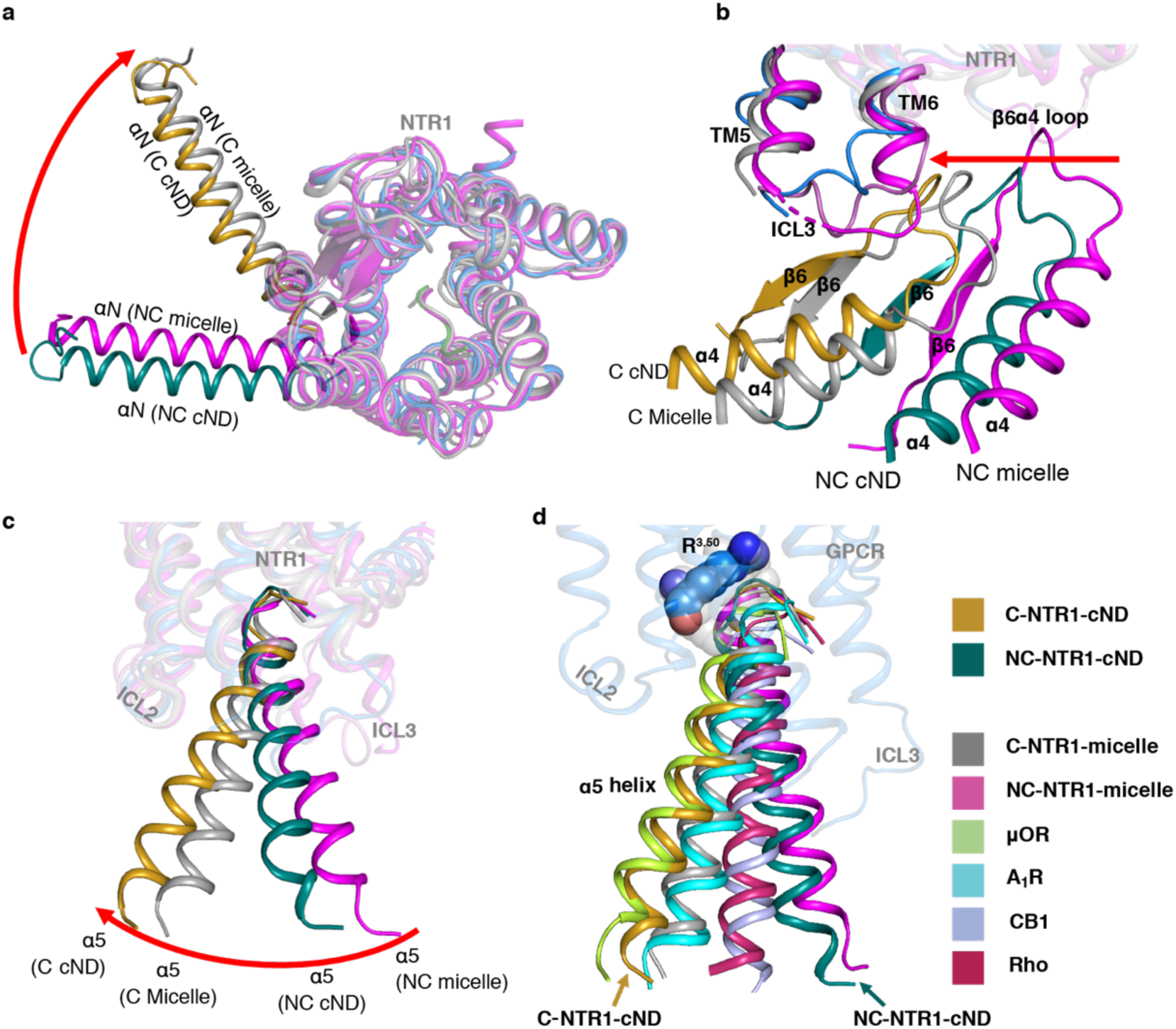
Comparison of NTR1-G_i_ interaction in lipid bilayer with detergent micelles. **a-c**, Superposed structure of C-state NTR1 (blue) and Gα (gold), NC-state NTR1 (orchid) and Gα (dark cyan), C-state NTR1 and Gα in micelle (gray), NC-state NTR1 and Gα in micelle (magenta). The models are superposed on NTR1. **a**, extracellular view of NTR1 and αN helix; **b**, side view of NTR1 ICL3 and β6α4 loop; **c**, side view of NTR1 and α5 helix. **d**, Comparison of the localization of α5 helix relative to GPCR in class A GPCR-G_i_ complex structures, including the canonical (gold) state and noncanonical (dark cyan) state structure reported in the current study, canonical (gray) and noncanonical (magenta) state of NTR1-G_i_ in detergent micelle, μOR-G_i_, Rho-G_i_, A_1_R-G_i_, and CB1-G_i_ in the same colors as panel (**b**). The structures are superposed on the GPCR. Residue R^3.50^ is shown as colored spheres in C-state NTR1 and as partially transparent gray spheres in the other GPCRs.

**Extended Data Fig. 11.**
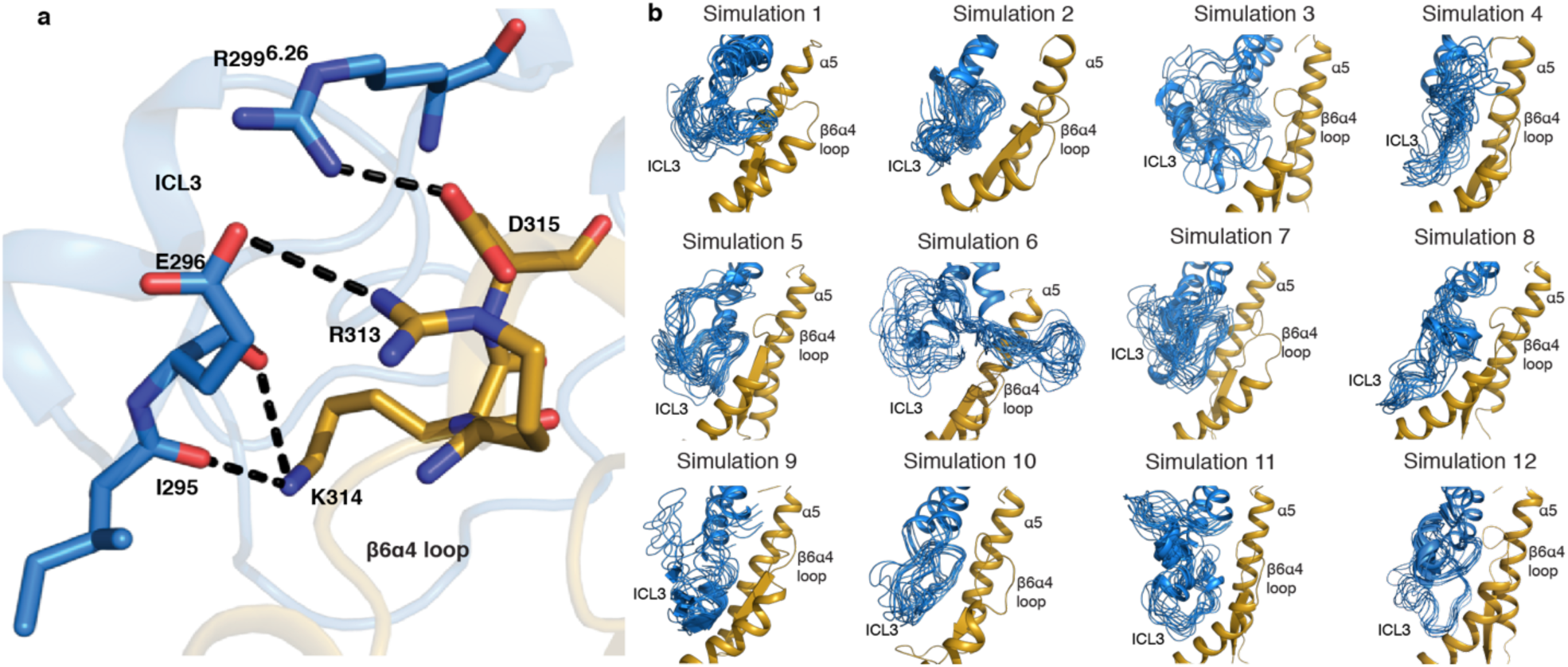
Molecular dynamics (MD) simulation for the interaction between ICL3 and the β6α4 loop. **a**, MD simulation showing salt bridges and hydrogen bonds form between TM6-ICL3 and β6α4-loop in the canonical state of NT-NTR1-Gi-cND represented by simulation 12. **b**, Dynamics of ICL3 for each independent simulation of the canonical state of NT-NTR1-G_i_-cND. Frames are sampled every 40 ns from each individual simulation. All 12 simulations show various interactions including salt bridges/hydrogen bonds between ICL3 and the β6α4-loop. An example of detailed interactions is shown in (**a**). NTR1 is colored in blue and G_i_ in gold in (**a-b**).

**Extended Data Table 1.** Cryo-EM data collection, refinement and validation statistics.

|  | NT-NTR1-G<sub>i</sub>-cND<br>Canonical state<br>(EMDB-xxxx)<br>(PDB xxxx) | NT-NTR1-G<sub>i</sub>-cND<br>Noncanonical state<br>(EMDB-xxxx)<br>(PDB xxxx) |
| --- | --- | --- |
| **Data collection and processing** |  |  |
| Magnification | 105,000 | 105,000 |
| Voltage (kV) | 300 | 300 |
| Electron exposure (e<sup>-</sup>/Å<sup>2</sup>) | 57 | 57 |
| Defocus range (μm) | -1.2 to -2.5 | -1.2 to -2.5 |
| Pixel size (Å) | 0.825 | 0.825 |
| Symmetry imposed | C1 | C1 |
| Initial particle images (no.) | 4,367,542 | 4,367,542 |
| Final particle images (no.) | 575,791 | 324,002 |
| Map resolution (Å) | 4.1 | 4.2 |
| FSC threshold | 0.143 | 0.143 |
| **Refinement** |  |  |
| Refinement software | Coot 0.9-pre, Phenix-dev-3318 |  |
| Initial model used (PDB code) | 4BUO, 1GP2, 6OY9 |  |
| Resolution limit set in refinement (Å) | 4.1 | 4.2 |
| Map sharpening *B* factor (Å<sup>2</sup>) | -245 | -204 |
| Model composition |  |  |
| Non-hydrogen atoms | 6959 | 6979 |
| Protein residues | 883 (6901 atoms) | 882 (6921 atoms) |
| Ligands | 1 (6 residues, 58 atoms) | 1 (6 residues, 58 atoms) |
| *B* factors (Å<sup>2</sup>) |  |  |
| Protein | 73.57 | 73.56 |
| Ligand | 66.76 | 66.76 |
| R.m.s. deviations |  |  |
| Bond lengths (Å) | 0.006 | 0.007 |
| Bond angles (°) | 1.052 | 1.365 |
| Validation |  |  |
| MolProbity score | 1.90 | 1.96 |
| Clashscore | 8.36 | 9.77 |
| EMRinger score | 1.46 | 1.17 |
| Poor rotamers (%) | 0.53 | 1.05 |
| Ramachandran plot |  |  |
| Favored (%) | 93.11 | 93.33 |
| Allowed (%) | 6.89 | 6.67 |
| Disallowed (%) | 0 | 0 |

## References

1. Griebel, G. & Holsboer, F. Neuropeptide receptor ligands as drugs for psychiatric diseases: The end of the beginning? Nature Reviews Drug Discovery 11, 462–478 (2012).

2. Shimada, I., Ueda, T., Kofuku, Y., Eddy, M. T. & Wüthrich, K. GPCR drug discovery: Integrating solution NMR data with crystal and cryo-EM structures. Nature Reviews Drug Discovery 18, 59–82 (2018).

3. Zhang, Y. et al. Cryo-EM structure of the activated GLP-1 receptor in complex with a G protein. Nature 546, 248–253 (2017).

4. Liang, Y. L. et al. Phase-plate cryo-EM structure of a class B GPCR-G-protein complex. Nature 546, 118–123 (2017).

5. Qi, X. et al. Cryo-EM structure of oxysterol-bound human Smoothened coupled to a heterotrimeric Gi. Nature (2019). doi: 10.1038/s41586-019-1286-0

6. Krishna Kumar, K. et al. Structure of a Signaling Cannabinoid Receptor 1-G Protein Complex. Cell 176, 448-458.e12 (2019).

7. Zhao, L.-H. et al. Structure and dynamics of the active human parathyroid hormone receptor-1. Science (80-.). 364, 148–153 (2019).

8. Kato, H. E. et al. Conformational transitions of a neurotensin receptor 1–Gi1 complex. Nature 572, 80–85 (2019).

9. Gao, Y. et al. Structures of the Rhodopsin-Transducin Complex: Insights into G-Protein Activation. Mol. Cell 75, 781-790.e3 (2019).

10. Tsai, C. J. et al. Crystal structure of rhodopsin in complex with a mini-Go sheds light on the principles of G protein selectivity. Sci. Adv. 4, (2018).

11. García-Nafría, J., Nehmé, R., Edwards, P. C. & Tate, C. G. Cryo-EM structure of the serotonin 5-HT1B receptor coupled to heterotrimeric Go. Nature 558, 620–623 (2018).

12. Draper-Joyce, C. J. et al. Structure of the adenosine-bound human adenosine A1 receptor-Gi complex. Nature 558, 559–563 (2018).

13. Liang, Y. L. et al. Cryo-EM structure of the active, G s -protein complexed, human CGRP receptor. Nature 561, 492–497 (2018).

14. Liang, Y. L. et al. Phase-plate cryo-EM structure of a biased agonistbound human GLP-1 receptor-Gs complex. Nature 555, 121–125 (2018).

15. Kang, Y. et al. Cryo-EM structure of human rhodopsin bound to an inhibitory G protein. Nature 558, 553–558 (2018).

16. Koehl, A. et al. Structure of the µ-opioid receptor-G i protein complex. Nature 558, 547–552 (2018).

17. García-Nafría, J., Lee, Y., Bai, X., Carpenter, B. & Tate, C. G. Cryo-EM structure of the adenosine A2A receptor coupled to an engineered heterotrimeric G protein. Elife 7, (2018).

18. Hilger, D., Masureel, M. & Kobilka, B. K. Structure and dynamics of GPCR signaling complexes. Nat. Struct. Mol. Biol. 25, 4–12 (2018).

19. Du, Y. et al. Assembly of a GPCR-G Protein Complex. Cell 177, (2019).

20. Rasmussen, S. G. F. et al. Crystal structure of the β 2 adrenergic receptor-Gs protein complex. Nature 477, 549–557 (2011).

21. Lee, A. G. How lipids affect the activities of integral membrane proteins. Biochimica et Biophysica Acta - Biomembranes 1666, 62–87 (2004).

22. Whorton, M. R. et al. Efficient coupling of transducin to monomeric rhodopsin in a phospholipid bilayer. J. Biol. Chem. 283, 4387–94 (2008).

23. Kofuku, Y. et al. Functional dynamics of deuterated β2-adrenergic receptor in lipid bilayers revealed by NMR spectroscopy. Angew. Chemie - Int. Ed. 53, 13376–13379 (2014).

24. Strohman, M. J. et al. Local membrane charge regulates β 2 adrenergic receptor coupling to G i3. Nat. Commun. 10, (2019).

25. Yen, H.-Y. et al. PtdIns(4,5)P2 stabilizes active states of GPCRs and enhances selectivity of G-protein coupling. Nature 559, 423–427 (2018).

26. Inagaki, S. et al. Modulation of the Interaction between Neurotensin Receptor NTS1 and Gq Protein by Lipid. J. Mol. Biol. 417, 95–111 (2012).

27. Kitabgi, P. Targeting neurotensin receptors with agonists and antagonists for therapeutic purposes. Curr. Opin. Drug Discov. Devel. 5, 764–76 (2002).

28. Nasr, M. L. et al. Covalently circularized nanodiscs for studying membrane proteins and viral entry. Nat. Methods 14, 49–52 (2016).

29. Wall, M. A. et al. The structure of the G protein heterotrimer Gi alpha 1 beta 1 gamma 2. Cell 83, 1047–58 (1995).

30. Egloff, P. et al. Structure of signaling-competent neurotensin receptor 1 obtained by directed evolution in *Escherichia coli*. Proc. Natl. Acad. Sci. 111, E655–E662 (2014).

31. Ballesteros, J. A. & Weinstein, H. Integrated methods for the construction of three-dimensional models and computational probing of structure-function relations in G protein-coupled receptors. Methods Neurosci. 25, 366–428 (1995).

32. Knepp, A. M., Grunbeck, A., Banerjee, S., Sakmar, T. P. & Huber, T. Direct Measurement of Thermal Stability of Expressed CCR5 and Stabilization by Small Molecule Ligands. Biochemistry 50, 502–511 (2011).

33. Krumm, B. E., White, J. F., Shah, P. & Grisshammer, R. Structural prerequisites for G-protein activation by the neurotensin receptor. Nat. Commun. 6, 7895 (2015).

34. Liu, X. et al. Structural Insights into the Process of GPCR-G Protein Complex Formation. Cell 177, 1243-1251.e12 (2019).

35. Dror, R. O. et al. Structural basis for nucleotide exchange in heterotrimeric G proteins. Science (80-.). 348, 1361–1365 (2015).

36. Sun, X., Singh, S., Blumer, K. J. & Bowman, G. R. Simulation of spontaneous G protein activation reveals a new intermediate driving GDP unbinding. Elife 7, (2018).

37. Chung, K. Y. et al. Conformational changes in the G protein Gs induced by the β 2 adrenergic receptor. Nature 477, 611–617 (2011).

38. Erlandson, S. C., McMahon, C. & Kruse, A. C. Structural Basis for G Protein–Coupled Receptor Signaling. Annu. Rev. Biophys. 47, 1–18 (2018).

39. Iiri, T., Bell, S. M., Baranski, T. J., Fujita, T. & Bourne, H. R. A Gsα mutant designed to inhibit receptor signaling through Gs. Proc. Natl. Acad. Sci. U. S. A. 96, 499–504 (1999).

40. Sun, D. et al. Probing Gα i1 protein activation at single-amino acid resolution. Nat. Struct. Mol. Biol. 22, 686–694 (2015).

41. Flock, T. et al. Universal allosteric mechanism for Gα activation by GPCRs. Nature 524, 173–179 (2015).

42. Goricanec, D. et al. Conformational dynamics of a G-protein α subunit is tightly regulated by Proc. Natl. Acad. Sci. 113, E3629–E3638 (2016).

## References

43. Egloff, P., Deluigi, M., Heine, P., Balada, S. & Plückthun, A. A cleavable ligand column for the rapid isolation of large quantities of homogeneous and functional neurotensin receptor 1 variants from E. coli. Protein Expr. Purif. 108, 106–114 (2015).

44. Hillenbrand, M., Schori, C., Schöppe, J. & Plückthun, A. Comprehensive analysis of heterotrimeric G-protein complex diversity and their interactions with GPCRs in solution. Proc. Natl. Acad. Sci. U. S. A. 112, E1181–90 (2015).

45. Delaglio, F. et al. NMRPipe: A multidimensional spectral processing system based on UNIX pipes. J. Biomol. NMR 6, 277–293 (1995).

46. Ehrenmann, J. et al. High-resolution crystal structure of parathyroid hormone 1 receptor in complex with a peptide agonist. Nat. Struct. Mol. Biol. 25, 1086–1092 (2018).

47. Schorb, M., Haberbosch, I., Hagen, W. J. H., Schwab, Y. & Mastronarde, D. N. Software tools for automated transmission electron microscopy. Nat. Methods 16, 471–477 (2019).

48. Zheng, S. Q. et al. MotionCor2: Anisotropic correction of beam-induced motion for improved cryo-electron microscopy. Nature Methods 14, 331–332 (2017).

49. Rohou, A. & Grigorieff, N. CTFFIND4: Fast and accurate defocus estimation from electron micrographs. J. Struct. Biol. 192, 216–221 (2015).

50. Tang, G. et al. EMAN2: An extensible image processing suite for electron microscopy. J. Struct. Biol. 157, 38–46 (2007).

51. Wagner, T. et al. SPHIRE-crYOLO is a fast and accurate fully automated particle picker for cryo-EM. Commun. Biol. 2, (2019).

52. Zivanov, J. et al. New tools for automated high-resolution cryo-EM structure determination in RELION-3. Elife 7, (2018).

53. Pettersen, E. F. et al. UCSF Chimera - A visualization system for exploratory research and analysis. J. Comput. Chem. 25, 1605–1612 (2004).

54. Emsley, P. & Cowtan, K. Coot: Model-building tools for molecular graphics. Acta Crystallogr. Sect. D Biol. Crystallogr. 60, 2126–2132 (2004).

55. Afonine, P. V. et al. Real-space refinement in PHENIX for cryo-EM and crystallography. Acta Crystallogr. Sect. D Struct. Biol. 74, 531–544 (2018).

56. Williams, C. J. et al. MolProbity: More and better reference data for improved all-atom structure validation. Protein Sci. 27, 293–315 (2018).

57. Epik, S. Schrödinger Suite 2018-2 Protein Preparation Wizard. (2019).

58. Madhavi Sastry, G., Adzhigirey, M., Day, T., Annabhimoju, R. & Sherman, W. Protein and ligand preparation: Parameters, protocols, and influence on virtual screening enrichments. J. Comput. Aided. Mol. Des. 27, 221–234 (2013).

59. Jo, S., Kim, T., Iyer, V. G. & Im, W. CHARMM-GUI: A web-based graphical user interface for CHARMM. J. Comput. Chem. 29, 1859–1865 (2008).

60. Lee, J. et al. CHARMM-GUI *Membrane Builder* for Complex Biological Membrane Simulations with Glycolipids and Lipoglycans. J. Chem. Theory Comput. 15, 775–786 (2019).

61. Lomize, M. A., Pogozheva, I. D., Joo, H., Mosberg, H. I. & Lomize, A. L. OPM database and PPM web server: Resources for positioning of proteins in membranes. Nucleic Acids Res. 40, (2012).

62. Case, D. A. et al. AMBER 2018. (2018).

63. Maier, J. A. et al. ff14SB: Improving the Accuracy of Protein Side Chain and Backbone Parameters from ff99SB. J. Chem. Theory Comput. 11, 3696–3713 (2015).

64. Dickson, C. J. et al. Lipid14: The amber lipid force field. J. Chem. Theory Comput. 10, 865–879 (2014).

65. Jorgensen, W. L., Chandrasekhar, J., Madura, J. D., Impey, R. W. & Klein, M. L. Comparison of simple potential functions for simulating liquid water. J. Chem. Phys. 79, 926–935 (1983).

66. Lee, J. et al. CHARMM-GUI Input Generator for NAMD, GROMACS, AMBER, OpenMM, and CHARMM/OpenMM Simulations Using the CHARMM36 Additive Force Field. J. Chem. Theory Comput. 12, 405–413 (2016).

